# Probing bacterial fitness in virtual chemostats

**DOI:** 10.64898/2026.10.31.763286

**Authors:** Kaan Öcal, Lars Maas, Michael P. H. Stumpf

**Affiliations:** School of BioSciences & School of Mathematics and Statistics, University of Melbourne, Parkville, Victoria 3052, Australia

**Keywords:** single-cell modelling, phenotypic variability, branching processes, particle filters

## Abstract

The growth rate of a bacterial population is a fundamental quantity in microbiology, but hard to predict from mechanistic single-cell models. Simulating exponentially growing cell populations is computationally infeasible, and single-lineage estimates are biased as they do not account for phenotypic selection between cells. We present computational algorithms to efficiently estimate growth rates with arbitrary accuracy by combining ideas from branching processes and particle filtering. These can be used with general agent-based cell models at a cost quadratic in the desired accuracy, bypassing the exponential complexity of naïve approaches. Applying these methods to single-cell models of *Escherichia coli*, we study how single-cell heterogeneity in antibiotic responses shapes population dynamics, and how bacterial locomotion and chemotaxis in fluctuating environments affect their long-term growth.

## 1 Introduction

The growth rate of a bacterial strain is one of the most important quantities in microbiology [1]: it is a major determinant of pathogenicity [2, 3], bacterial yield in biotechnology [4–6] and evolutionary fitness [7–16]. Despite significant strides in large-scale single-cell modelling [17–21], our ability to predict population growth rates is remarkably limited, particularly in the presence of cell-to-cell variability [3, 22, 23]. Bacterial populations are typically too large to simulate individually, and single-lineage estimates often give incorrect answers as they neglect the intricate interplay between fitness, selection and individual variability in real populations [24–27]. In this paper, we present algorithms for agent-based populations (“virtual chemostats”) that can efficiently predict population quantities such as growth rates to arbitrary accuracy.

Our approach uses ideas from Sequential Monte Carlo and particle filtering [28, 29] to randomly subsample cells in a population, and is closely related to cloning algorithms in statistical physics [30–32]. The resulting algorithms can estimate population quantities with a computational effort that scales quadratically with the desired accuracy: halving the statistical error requires four times the computational effort, in line with most Monte Carlo approaches, and unlike the exponential cost of simulating a full population. Importantly, tracking isolated lineages as in a mother machine [33] systematically underestimates population growth rates, and does not give the correct answer even in the long run [27]. We show that this phenomenon, which is known as nonlinearity bias in the statistical physics literature, arises from the failure of single-lineage experiments to account for phenotypic selection in a population [34]. By expanding the population to allow multiple competing lineages, we can reintroduce these missing selective forces and obtain accurate estimates of population dynamics and growth rates from relatively few (on the order of 20–100) lineages.

The ideas presented in this paper build on recent connections between microbial population behaviour and the statistical physics of lineages [25, 35–38]. While mathematical formulæ for growth rates can be derived e.g. in the form of the generalised Euler-Lotka equation [38– 40], for biophysically realistic models these are typically infinite-dimensional eigenvalue equations and numerically intractable, with exact solutions only available in limited cases [39, 41–44]. Computing growth rates in fluctuating environments is particularly difficult, and analytical results have remained sparse despite intense interest [23, 45–48]. We showcase the potential of our numerical approach by studying how the stochastic response of the *tet*-operon in *Escherichia coli* shapes a population’s ability to survive antibiotic exposure, and by investigating how motile bacteria use chemotaxis to maximise nutrient intake and long-term growth in an unpredictable environment.

## 2 Theory

### 2.1 Estimating growth rates

We view cells as individuals that grow and either die or divide into two daughter cells. Mathematically, this describes a branching process (see App. A). For simplicity we do not consider any direct interactions between cells, such as density effects or resource depletion. We furthermore assume that all cells form one strain and neglect long-term effects such as genetic mutations. Under these assumptions, the number of cells in a population will grow roughly exponentially in time:

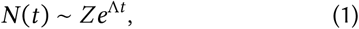

where Λ is the population growth rate, which is an intrinsic property of the strain in a given environment, and *Z* is a (random) prefactor that incorporates differences in initial colony size, lag phase effects and single-cell variability.

We visualise the growth curves predicted by Eq. (1) in Fig. 1**a**. As expected, log *N* (*t*) forms a straight line for large *t*, with fixed slope Λ and a random intercept equal to log *Z* that varies between runs. As long as the population does not go extinct, we can formally recover the growth rate as

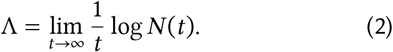

**Figure 1.**
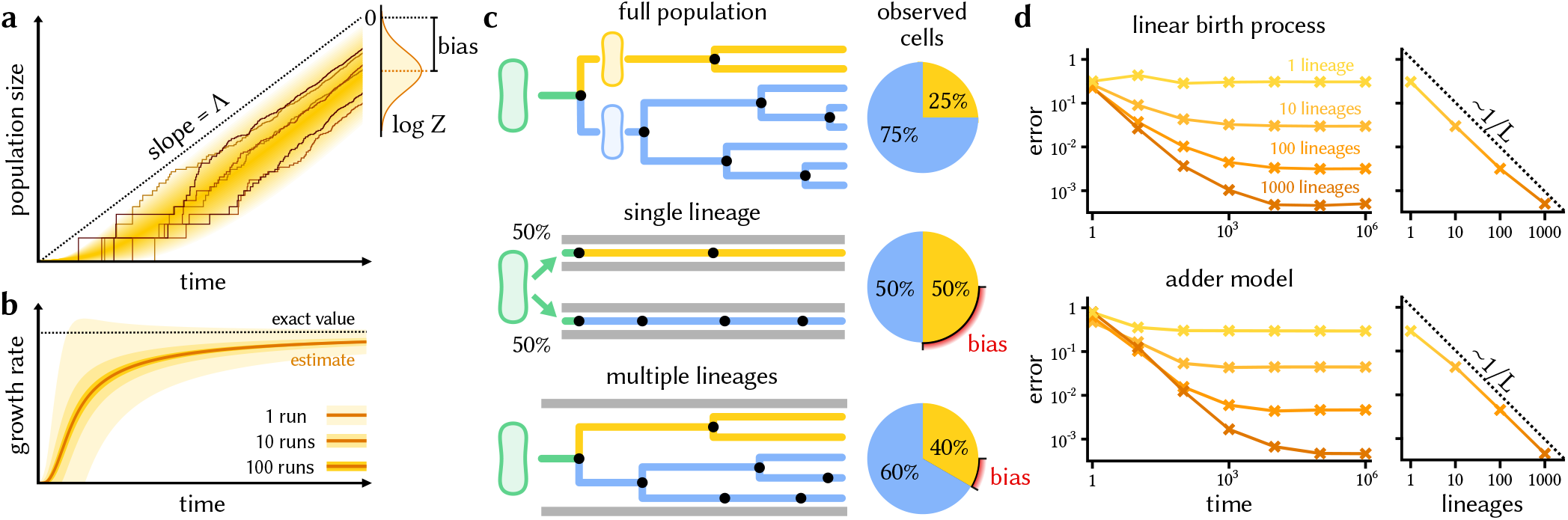
Estimating growth in bacterial populations. **a** Long-term behaviour of a freely growing population, with population size plotted on a log scale. Solid lines show individual realisations of the population process starting from one cell, and the shaded area indicates average behaviour. For each run, the log population size will eventually have slope Λ (dotted line for reference). The histogram outset shows the distribution of the intercept log *Z*, which affects the growth rate estimator 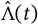). If log *Z* has nonzero average, estimates will be systematically biased. **b** Behaviour of the estimator 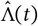 in Eq. (3) for the same population process. The solid line indicates the estimator average, the shaded area indicates the standard error for different sample sizes, and the dotted line represents the true growth rate. Averaging multiple estimates brings the result closer to the mean, which differs from the true growth rate due to finite-time bias. **c** A mother cell dividing into two unequal daughter cells, with the upper daughter reproducing more slowly than the lower daughter. Most cells in the population (top) descend from the fitter daughter. When tracking a single lineage, as in an idealised mother machine, we observe the slower daughter and its descendants in 50% of cases (middle). The effects of selection are invisible, and we will underestimate the population growth rate on average. In a chemostat where multiple lineages can coexist (bottom), some amount of selection can take place, leading to improved growth estimates. **d** Root mean square error of the growth rate estimate as a function of time and the number of lineages tracked, for two simple models of cell growth with true growth rate Λ = 1. The single-lineage case corresponds to Eq. (6). For a fixed population size *L*, the finite-size error is of order *O*(1/*L*). Lineages were simulated in a strict chemostat; see App. H for details on the two models.

In practice, we can only observe a bacterial population for a finite time *t*, which leads to the finite-time estimator

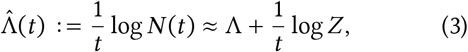

where we assume *N* (0) = 1. The intercept log *Z* in Fig. 1**a**, which represents random variation in population size, distorts our estimate 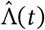 by an amount asymptotically proportional to 1/*t*. Following [27], we call this *finite-time bias*. In theory, as *t* → ∞, this bias becomes less important and our estimate converges to the true growth rate Λ; in practice, the number of cells grows exponentially in *t*, which limits the accuracy we can hope to obtain.

Instead of simulating an exponentially growing number of cells, we could attempt to reduce the error by averaging multiple smaller populations. This does not eliminate our bias, however. Indeed, assume we have estimates 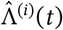 for *M* independent realisations of our population for fixed *t*. Averaging these estimates will decrease sampling noise by a factor of 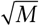, and one might hope to obtain something close to Λ given enough samples. Instead, for large *M* we find

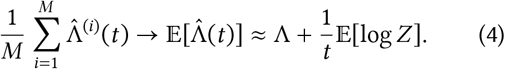

Since E[log *Z*] ≠ 0 in general, the finite-time bias in Eq. (3) does not disappear upon simple averaging, as shown in Fig. 1**b**. Attempting to eliminate this bias by starting with a larger initial population, or pooling populations, runs into similar difficulties (see App. B).

This finite-time error in estimating Λ is to be expected [27]: we cannot measure the long-term growth rate of a population from only short-time observations. For example, *E. coli* can divide every 20 minutes under ideal conditions [49], and we cannot hope to get a good estimate of this number in less than 20 minutes. In practice, it may take many generations for us to accurately infer Λ, especially in the presence of large-scale environmental fluctuations.

The most reliable way to decrease the finite-time bias in our estimates of Λ is to increase *t*. The difficulty of doing so in an exponentially growing population is immediate: going from 10 to 20 generations, for example, halves the error, but requires 2^10^ ≈ 1000 times as many cells. Ideally, we would need only 4 times as many samples for the same improvement (averaging *M* samples should decrease sampling error by a factor of 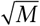). To accurately extract growth rates from finite data, we must therefore find better statistical estimators that can probe the long-term dynamics of a population while bypassing this exponential scaling.

#### Single-lineage tracking

To understand the behaviour of a population, we do not need to track every single cell. Instead, we can extrapolate population behaviour and growth from individual lineages [25]. As we review in App. C and Fig. S1**a**, the total size of our population is related to the number of divisions *n*(*t*) in a typical lineage as follows:

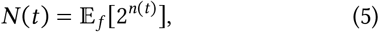

neglecting cell death for simplicity. Here we average using the forward distribution over lineages, where we start with the original ancestor and pick a random daughter cell at each division. This matches the behaviour of an idealised mother machine [25]. Eq. (5) suggests that we can estimate *N* (*t*), and therefore Λ, by subsampling *L* ≪ *N* (*t*) random lineages, with statistical error depending on the number *L* of sampled lineages. As we shall see, however, results depend strongly on how this is done.

The standard approach is to track a single lineage, for example in a mother machine, and to count the number of divisions. Assuming an ideal mother machine where each daughter has a 50% probability of being chosen, this yields the familiar single-lineage estimator for the growth rate:

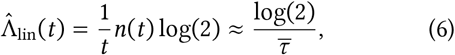

where 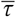 is the mean generation length in the lineage. This, however, does not yield good estimates in general. Even for *t* → ∞, Eq. (6) does not converge to Λ: the single-lineage estimator is not statistically consistent [27]. Although this seems counterintuitive, the error can be quite substantial. For example, if all cells have independent exponentially distributed lifetimes, one can calculate that the relative error will be on the order of 30% (App. G).

The reason for this discrepancy is that tracking a single lineage does not take into account selection. Wherever there is variability within a population, cells that divide faster will contribute more offspring in the future; as a result, the population will grow faster than expected by looking at single lineages [34]. This is illustrated in Fig. 1**c**, where we show a cell dividing into two unequal daughter cells, one of which divides faster than the other. In the long run, more than half of the population will descend from the fitter cell; in contrast, when tracking a single lineage, half the time we will extrapolate from the slower daughter. Since this scenario occurs with every new division, the single-lineage estimate will consistently underestimate Λ even as *t* → ∞. We numerically verify this behaviour for two exactly solvable models in Fig. 1**d**, the linear birth process and the adder model; the error of the single-lineage estimate remains above 10% even after a million generations.

We call this error *finite-size bias*, as it arises when we extrapolate population behaviour from a small number of lineages [27]. Unlike finite-time bias, which also occurs in single-lineage estimates, finite-size bias does not disappear when *t* → ∞. It can be reduced by exponentially averaging Eq. (6) as suggested by Eq. (5), but the number of lineages required to overcome this bias grows exponentially in *t* [27, 50, 51]. In this sense, the single-lineage estimator is not much more efficient than the naïve approach Eq. (3).

This new form of bias is known as nonlinearity bias in statistical physics [27, 50, 51]. In a biological context, it arises as a systematic error from neglecting phenotypic selection. Finite-size bias increases when there is more cell-to-cell variability, as this increases the amount of selection. Only when all cells divide at exactly the same time does Eq. (6) yield the correct long-time estimate.

While Eq. (6) is not a good estimator for the growth rate Λ in the presence of cell variability, it can be used to quantify the amount of selection inside a population [37]. Phenotypic selection in a population favours fitter cells, and as a result increases growth rates. In the absence of cell death, this growth rate gain equals the Kullback-Leibler divergence rate between the forward and backward distributions over lineages:

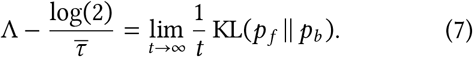

As before, the forward distribution *p*_*f*_ represents the statistics of a lineage observed by tracking the descendants of a single cell (for example in a mother machine), whereas the backward distribution *p*_*b*_ represents the ancestral lineage of a random cell in the full population (see App. C). Unlike the forward distribution, the backward distribution is shaped by selection, since a cell that has more descendants will appear in more ancestral lineages. The Kullback-Leibler divergence rate is zero precisely if forward and backward statistics are the same, *p*_*f*_ = *p*_*b*_, which indicates that all lineages are equally fit. Otherwise, the difference between *p*_*f*_ and *p*_*b*_ indicates (phenotypic) traits subject to selection, the effect of which is to increase the population growth rate beyond the naïve average suggested by Eq. (6) [25, 37].

Of course, a fundamental shortcoming of the estimator Eq. (6) is that it only works for surviving lineages, and it is not at all clear how to interpret it when a cell dies. Since cell death is just a form of selection, this underlines the inability of single-lineage measurements to deal with selective forces. For some models of cell growth, single-lineage measurements can be analytically corrected to account for the resulting bias [52–55], but deriving these corrections is difficult in general.

### 2.2 Population control

We have discussed two very different ways of estimating growth rates. The direct approach observes the entire population, but suffers from substantial finite-time bias in practice. The single-lineage estimator Eq. (6) only tracks a single cell at a time, which reduces the severity of this finite-time bias, at the cost of introducing an additional finite-size bias. Both approaches require an exponential amount of computational power to overcome their respective biases. In the following, we present estimators that interpolate between simulating the entire population and simulating a single lineage, therefore trading finite-time bias for finite-size bias. By doing this systematically, we arrive at algorithms that eliminate the exponential over-head of the previous two approaches, and instead have quadratic computational cost in the desired accuracy.

#### Thinning

Returning to Eq. (5), a simple way to subsample lineages is to introduce an artificial death rate *δ* > 0 into the population. Here, cells are removed at random from the population with exponential rate *δ*, simulating a virtual chemostat with fixed outflow rate. The probability that a lineage from the original population survives until time *t* is *e*^−*δt*^, so the observed population size *N*_obs_ is related to the true population size as

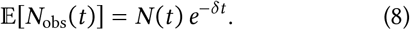

The observed population grows exponentially with rate Λ − *δ*, and we can estimate the growth rate of the full population as

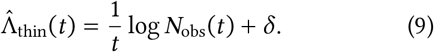

We visualise this approach in Fig. 2**b**.

**Figure 2.**
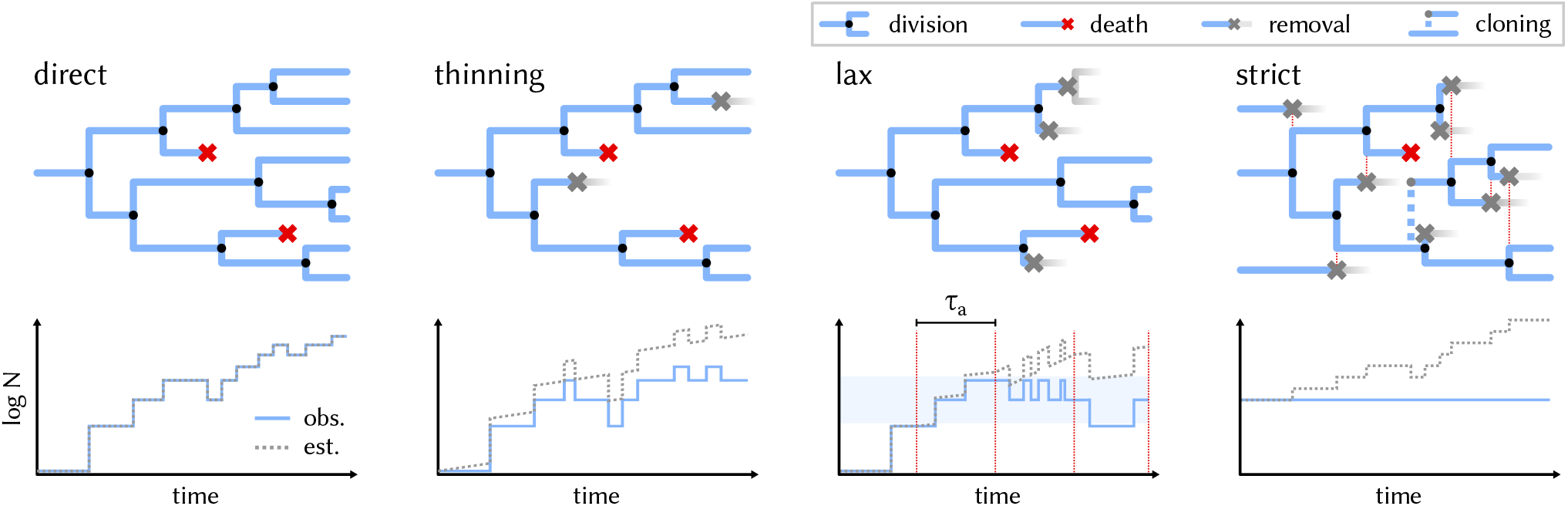
Statistical algorithms to simulate growing populations. The direct approach simulates all cells in a population, with exponentially increasing computational cost. Thinning removes cells at a fixed rate *δ*, which reduces the speed of population growth, but not its exponential nature. The true population size is estimated via Eq. (8). The lax chemostat adapts the thinning rate to the current growth rate at regular intervals *τ*_*a*_ to maintain the population size around *L* (shaded area). The true population size is estimated via Eq. (10). The strict chemostat (a variant of the cloning algorithm [30, 31]) keeps a fixed population size *L* by discarding one cell at random from the population at each division, and cloning a random cell whenever an existing cell dies. The true population size is estimated via Eq. (16). This requires synchronising cell division and death events (dotted lines).

One disadvantage of this approach is that introducing *δ* means that cells are more likely to die, leading to the risk of extinction for the whole population. In fact, for *δ* ≥ Λ, the thinned population has negative or zero growth rate, and extinction is certain in the long run. For *δ* < Λ, thinning will simulate an exponentially growing population, albeit at a reduced rate compared to the original. Tuning *δ* involves a trade-off between decreasing the simulated growth rate and increasing the chance of extinction. Of course, since Λ is unknown *a priori*, this is not straightforward in practice.

No matter how *δ* is chosen, however, thinning has one distinguishing feature: every lineage has the same probability of being removed. In this sense, thinning returns an unbiased subset of the true population, cf. Eq. (8). This does not mean that the resulting growth rate estimate is unbiased; as we discussed earlier, this is not even the case for the full population. However, as we will see below, unbiased sampling is the starting point for other subsampling schemes that allow us to estimate Λ with low bias, and without the exponential cost of the direct (or thinned) approach.

#### The lax chemostat

One way to improve thinning is to allow *δ* to vary as a function of *t*, depending on the observed population size. For example, we can set *δ*(*t*) ≈ 0 when the population size is small to minimise the risk of extinction, and increase *δ*(*t*) when the population size becomes too large. We can therefore maintain the population size around any target value *L* by periodically adapting *δ*. We call this method the lax chemostat, as it allows the “outflow” rate, and therefore the population size, to fluctuate with time (strictly speaking, this describes a turbidostat). This resembles the approach used in [56], where cells are subsampled when the population size reaches a certain threshold. For the lax chemostat, we can estimate the full population size as

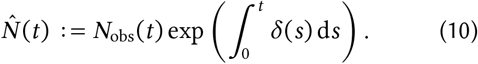

In App. D, we show that this is an unbiased estimator of *N* (*t*). We also obtain the growth rate estimate

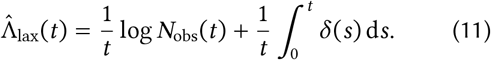

Unlike Eq. (10), the growth rate estimate 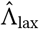 is biased: taking the logarithm of the unbiased estimator in Eq. (10) will systematically underestimate Λ, as one can show using Jensen’s inequality. This bias arises from the fact that we only track a subset of the population to estimate *N* (*t*), and is another form of finite-size bias. Unlike the single-lineage case, however, this finite-size bias is on the order of *O*(1/*L*), depending on the target population size *L* [32]. We can thus reduce the bias by increasing *L*, at the cost of simulating more lineages. Taking into account the effects of finite-time bias, finite-size bias and sampling variance, in App. D we argue that the root mean square error (RMSE) of the lax chemostat is asymptotically

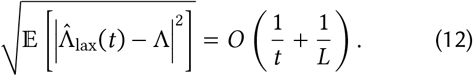

Since the computational complexity of the lax chemostat is *O*(*Lt*), this implies that our estimator is relatively efficient: we need quadratically many samples to obtain linear accuracy in our estimates, as measured by the root mean square error (RMSE) [57]. In fact, we can halve our estimation error by doubling both *L* and *t*, which will quadruple the computational cost. This is quite unlike the exponential complexity of the earlier approaches, and in line with the quadratic scaling of standard Monte Carlo estimators.

In practical terms, the lax chemostat with a target population size *L* periodically updates *δ*(*t*) at intervals of length *τ*_*a*_, a user-defined adaptation time. To find *δ*(*t*), we estimate the growth rate in the previous interval as

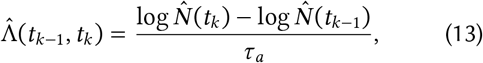

where *t*_*k*_ = *kτ*_*a*_ is the *k*-th timepoint. We then set the death rate for the next interval by requiring that

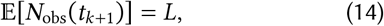

which combined with Eq. (13) yields

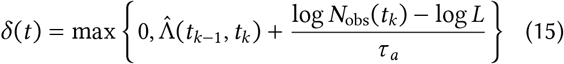

for *t*_*k*_ ≤ *t* < *t*_*k*+1_. Intuitively, Eq. (15) adds a variable death rate that steers the population size towards *L*. If *N*_obs_(*t*_*k*_) is below a critical threshold, we set *δ*(*t*) = 0 to avoid the risk of extinction. This algorithm is visualised in Fig. 2**c**; we provide pseudocode in Algorithm 1 in the Appendix. A major advantage of this approach is that cells in the lax chemostat evolve independently within each update interval (*t*_*k*_, *t*_*k*+1_), and can be simulated in parallel. A disadvantage is that fluctuations in the population size can still drive the population to extinction, which can be counteracted in two ways. We can either reduce the adaptation time *τ*_*a*_, allowing *δ*(*t*) to quickly react to low population sizes, or increase *L*, which suppresses the probability of extinction exponentially. In our applications, we found the lax chemostat to work reliably for *L* ≥ 50 if the adaptation time roughly matches the population doubling time. In practice, this will depend on the problem; we can also replace Eq. (13) with a running average to obtain more stable estimates. For more details on the implementation of the lax chemostat, see App. F.

#### The strict chemostat

The lax chemostat resembles a well-known class of methods in statistical physics, commonly called cloning algorithms [30, 31]. Inspired by these, we can define a “strict” chemostat by requiring that the population size is kept constant at a fixed value *L* at all times, see e.g. [58]. When a cell divides in two, we discard one of the *L* + 1 resulting cells at random, and when a cell dies, we copy (“clone”) a random other cell in its place.

This approach is reminiscent of the Moran process in population dynamics. When *L* = 1, it is equivalent to tracking a single lineage as discussed above. We visualise this algorithm in Fig. 2**d**, and provide pseudocode in Algorithm 2 in the Appendix.

For the strict chemostat, we can estimate the true population size via

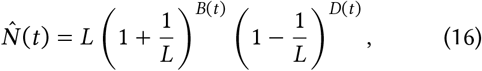

where *B*(*t*) is the number of division (or birth) events and *D*(*t*) the number of deaths up to time *t*, as in [6, 59]. At each division, we track *L* out of the *L*+ 1 resulting bacteria at random, so our observed population underestimates the later population size by a factor of *L*/(*L*+1) on average. Similarly, at each death we artificially inflate the population size by 1 to reach *L*, therefore overestimating the later population size by a factor of *L*/(*L* − 1). We show that Eq. (16) yields an unbiased estimate of E[*N* (*t*)] in App. E. Note that setting *L* = 1 and neglecting cell death (which would result in extinction) precisely recovers Eq. (5). From Eq. (16) we obtain the growth rate estimate

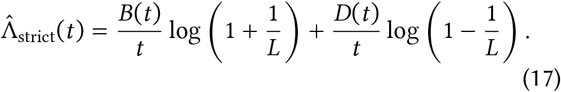

For statistical correctness, the cell to be discarded following a division event must be chosen at random from the whole population, and not just among the two daughter cells. Indeed, discarding a daughter cell at random is equivalent to simulating multiple independent lineages [60], which does not yield accurate growth rate estimates as we saw earlier. Our prescription allows lineages to compete for space in the chemostat, which captures the effects of selection and results in better estimates. We analyse the asymptotic behaviour of the strict chemostat in App. E, based on [32], and deduce the same asymptotic efficiency as its lax counterpart. The predicted finite-size bias of order *O*(1/*L*) matches our numerical results in Fig. 1**d**. The single-lineage algorithm with *L* = 1 incurs the largest bias, as it captures the least amount of selection.

A disadvantage of the strict chemostat is that it is not easily parallelised. Indeed, our prescription requires that we synchronise all lineages whenever a cell dies or divides (see Fig. 2**d**). This makes this approach less efficient for large *L*. Unlike the lax chemostat, the strict chemostat cannot go extinct for *L* > 1, as cells are only removed when there is a surplus, and dead cells are replaced by clones of live ones. Instead, the strict chemostat can stall when all cells are close to death, as it will keep cloning cells that die immediately afterwards. To prevent this and similar difficulties associated with cloning, we describe a modified version of this algorithm in App. E that does not clone cells upon death, thus allowing the population size to drop below *L*.

The strict and the lax chemostat algorithms are fundamentally based on the same principle: while the population size is controlled by continually removing cells, each cell has the same probability of being removed at each step. We therefore take unbiased subsamples of the population. By keeping track of how many cells are removed, we can extrapolate the simulated population to estimate the size of the original population, and therefore its growth rate. Viewed from this perspective, both algorithms are effectively particle filters [28, 29] and represent a form of Sequential Monte Carlo, which explains their favourable statistical properties in the long run. Importantly, both approaches are statistically far more efficient than simulating multiple independent lineages.

As we show in App. E for the strict chemostat, this form of population control allows us to compute more general population averages with similar accuracy to growth rates [59]. Furthermore, by tracing the ancestry of a random cell we obtain approximate backward lineages for large *t* [6], which allows us to probe the effects of phenotypic selection inside a population. We remark that a single population effectively only yields one backward lineage in the long run, even for large *L*, since coalescence means that all cells will share the same ancestor around *L* generations ago on average.

We recapitulate the statistical and computational properties of the estimators discussed here in Table 1. In general, we recommend the strict chemostat for small population sizes on the order of *L* = 1–25 due to its stability, and the lax chemostat for *L* ≳ 25 (where extinction is less of a concern) due to its parallelisability. Implementation details for both algorithms can be found in App. F.

**Table 1.**
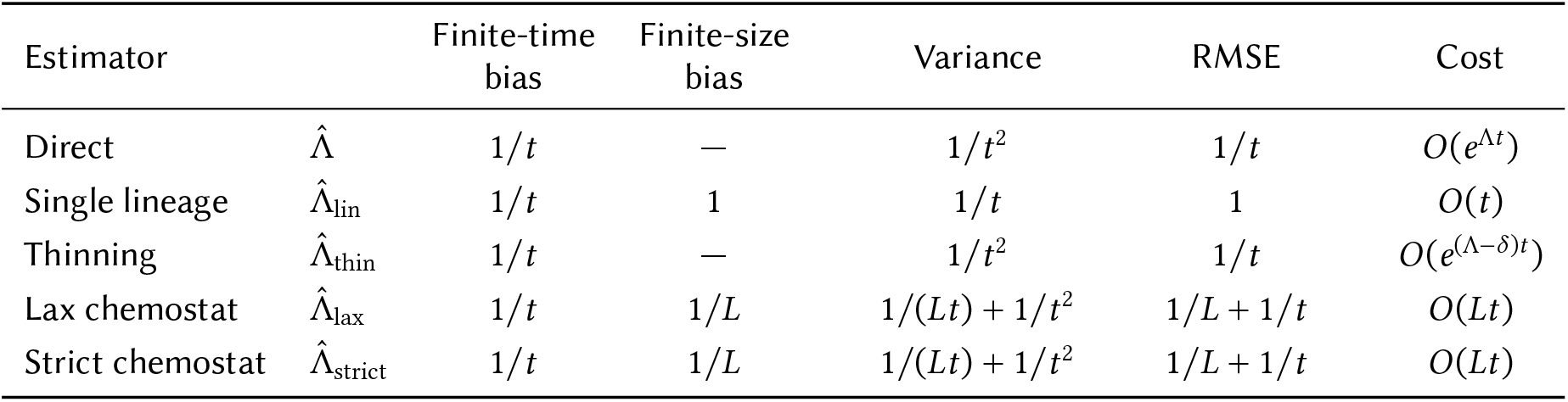
Comparison of the growth rate estimators discussed in this paper. Entries show how each quantity scales with *t* and *L*, up to constant factors. Computational cost is measured in the number of cells simulated.

| Estimator |  | Finite-time bias | Finite-size bias | Variance | RMSE | Cost |
|---|---|---|---|---|---|---|
| Direct | Λ̂ | 1/t | — | 1/t<sup>2</sup> | 1/t | O(e<sup>Λt</sup>) |
| Single lineage | Λ̂<sub>lin</sub> | 1/t | 1 | 1/t | 1 | O(t) |
| Thinning | Λ̂<sub>thin</sub> | 1/t | — | 1/t<sup>2</sup> | 1/t | O(e<sup>(Λ−δ)t</sup>) |
| Lax chemostat | Λ̂<sub>lax</sub> | 1/t | 1/L | 1/(Lt) + 1/t<sup>2</sup> | 1/L + 1/t | O(Lt) |
| Strict chemostat | Λ̂<sub>strict</sub> | 1/t | 1/L | 1/(Lt) + 1/t<sup>2</sup> | 1/L + 1/t | O(Lt) |

## 3 Applications

### 3.1 Tetracycline response in *E. coli*

We apply our results to study how the antibiotic response mechanism encoded by the *tet*-operon affects population growth and survival in *E. coli* [61]. The *tet*-operon (Fig. 3**a**) is a widespread form of antibiotic resistance mediated by the efflux protein TetA, which actively removes antibiotics from the cytosol. Under normal conditions, TetA is transcriptionally repressed by TetR, a dimeric transcription factor that is encoded in close proximity to *tetA*. TetR binds to two *tetO* operator sequences that overlap the transcription start sites of both *tetA* and *tetR*, leading to a negative feedback loop. When tetracycline analogues enter the cell, they bind allosterically to TetR, causing the latter to dissociate from the DNA. This in turn results in the production of TetA to export the antibiotic from the cell.

**Figure 3.**
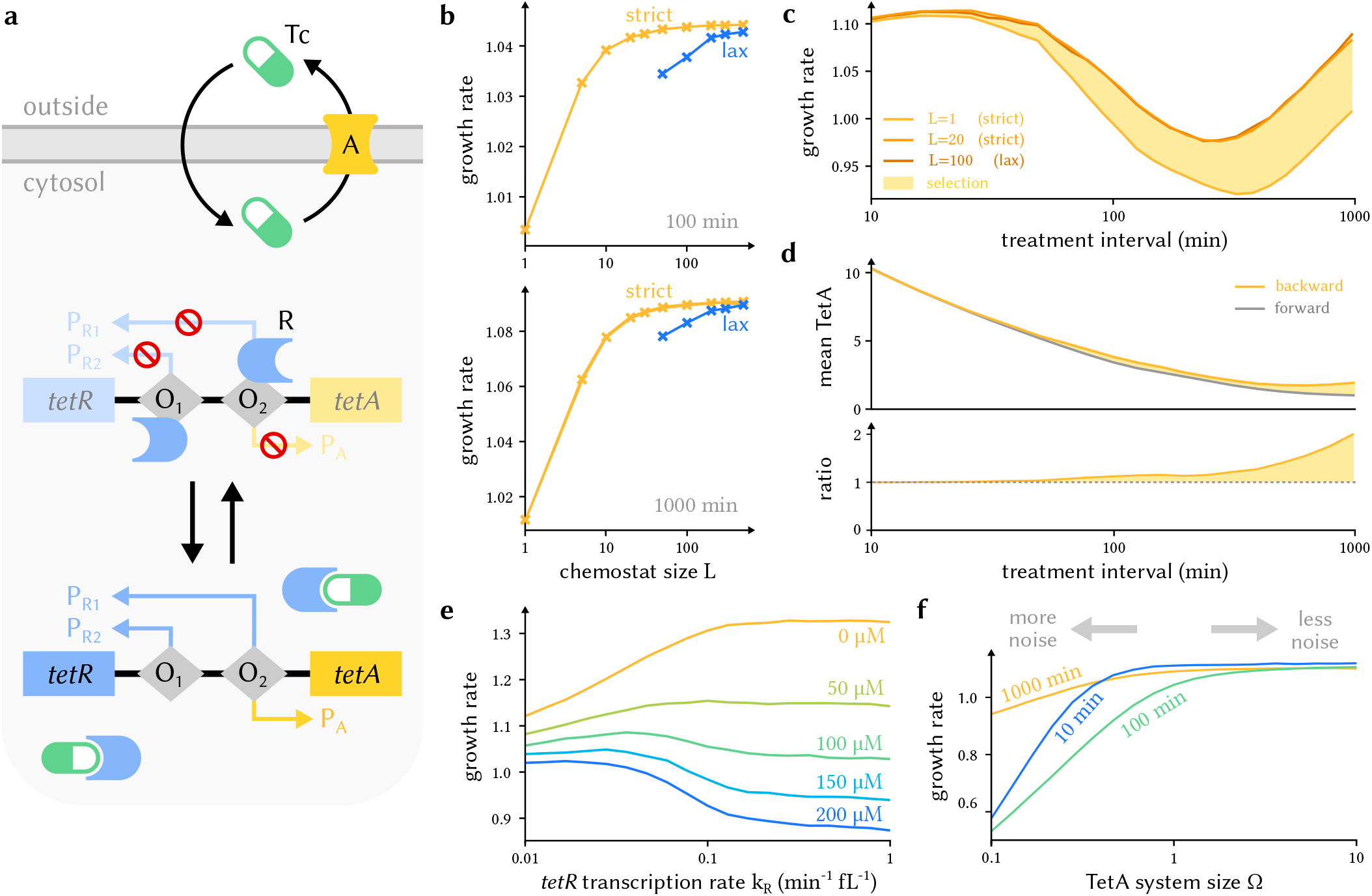
Tetracycline resistance encoded by the *tet*-operon. **a** Illustration of the model. Tetracycline (Tc) can diffuse into the cell and is actively exported by TetA (A). The TetR repressor (R) is normally bound to the *tet*-operon and blocks transcription of both *tetR* and *tetA*. When tetracycline binds to TetR, TetR dissociates from the operon, allowing transcription of both genes. The operon includes two TetR-binding sites O_1_ and O_2_ that overlap the transcription start sites of both genes, blocking transcription when bound. TetR is still produced at a reduced rate when O_2_ is bound, suggesting the presence of a second transcription start site not overlapping O_2_. **b** Growth rate estimates depending on the simulated population size, for both the strict and the lax chemostat. As the population size increases, the effects of selection are captured more accurately, and the growth estimate converges to a fixed value. The lax chemostat is less accurate for fixed *L*, since the population size can fluctuate below *L*. The single-lineage estimate (*L* = 1) is noticeably below the likely true value. **c** Population growth rate as a function of the treatment interval, for a fixed total dose. The estimates for *L* = 20 and *L* = 100 are very similar and indicate convergence (see **b**). The difference between the population and the single-lineage estimate (shaded area) measures the selection-induced growth rate gain. When tetracycline is applied rapidly, cells maintain stable TetA levels and selection is weak; when tetracycline is applied infrequently, single-cell variability in TetA expression at each exposure event leads to increased selection. **d** Mean TetA levels measured in forward and backward lineages, and the ratio between the two. When tetracycline exposure is infrequent, ancestral cells (as measured by backward lineages) express on average twice as much TetA as typical cells in a forward lineage; this difference leads to phenotypic selection between cells. **e** Growth rate at different tetracycline concentrations as a function of TetR expression, with fixed treatment interval of 100 minutes. At low TetR levels, TetA is overproduced and inhibits growth in the absence of tetracycline, but protects the cell at high doses; at high TetR levels, cells grow faster under normal conditions, but become more susceptible to tetracycline. The transcription rate *k*_R_ trades off robustness against maximal growth rate. **f** Growth rate as a function of molecular noise due to TetA expression, measured by the system size parameter Ω. At low Ω, the tetracycline response becomes noisy and inhibits growth, but the effect levels off around Ω = 1, corresponding to the average TetA levels measured in **d.** All growth rates are given in doublings per hour. Unless otherwise noted, parameters are given in Table S1. Further model and simulation details can be found in App. I.

The *tet*-operon encodes an adaptive response, as TetA is only produced in the presence of tetracycline owing to its tight repression by TetR [62]. This is necessary since TetA reduces the membrane potential and is toxic to normally growing cells [63]. Although tetracycline binds strongly to TetR, leading to a very sensitive response, outcomes are highly heterogeneous between cells. Many cells fail to produce enough TetA before their translational machinery is incapacitated by tetracycline, and enter a prolonged state of arrest until either tetracycline exposure stops or enough TetA is produced to decrease tetracycline concentrations sufficiently. Other cells overproduce TetA owing to insufficient TetR expression, and enter a moribund state due to TetA toxicity [64]. This variability in outcomes, where only a fraction of cells manage to quickly overcome tetra-cycline exposure, can be explained by the low number of TetR and TetA molecules in cells [65], resulting in significant gene expression noise that dramatically affects cell survival. This raises the question why this response has not evolved to become less noisy, e.g. by increasing TetR and TetA expression levels.

To study the population-level impact of this variability in tetracycline response, we use a stochastic model of the *tet*-operon based on [65]. Following the metabolic model of [66], cells grow exponentially by accumulating biomass largely due to protein production, with a rate λ that is determined by the translational capacity κ_*t*_ and the excess fraction of cellular resources allocated to ribosomes, 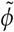:

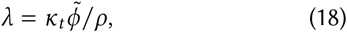

where ρ is a proportionality factor. The excess fraction 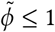 is determined by the nutritional capacity κ_*n*_ as

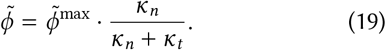

This reflects a trade-off due to resource division between ribosome-affiliated proteins, which directly contribute to cell growth, and metabolic proteins required for normal cell function [66]. We incorporate the ribosome-inhibiting effect of tetracycline and TetA toxicity into the translational capacity via non-competitive inhibition:

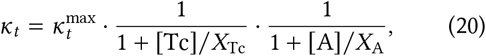

where 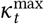 is the maximal translational capacity, [Tc] and [A] denote tetracycline and TetA concentrations, respectively, and *X*_Tc_ and *X*_A_ are the inhibition constants, i.e. the concentrations of tetracycline and TetA at which the translational capacity κ_*t*_ is halved.

We model stochastic production of TetR and TetA as in [65]. For simplicity, we do not consider transcription and translation separately, combining them into a single production step occurring with propensities

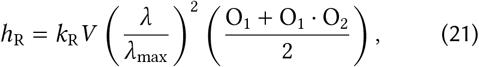

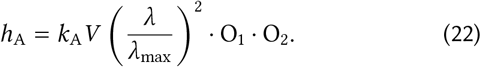

Here each indicator variable O_1_ and O_2_ is 1 when the site is free, and 0 when it is blocked by TetR (we model these probabilistically using the quasiequilibrium approximation, see App. I). The growth-rate-dependent term arises since both transcription and translation scale with λ, where λ_max_ is the maximal growth rate possible under Eq. (18) and Eq. (20). Eq. (21) reflects the fact that TetR has two transcription start sites, and can be transcribed at a reduced rate even when O_2_ is bound.

Due to the extremely tight dimerisation of TetR [67], we model TetR as an obligate dimer with at most one unmatched molecule at any given time. Tetracycline binds to each TetR dimer with dissociation constant *K*_RTc_, which we model using the quasiequilibrium approximation owing to the rapid speed of binding and unbinding [67]. We similarly model the binding of free TetR dimers to O_1_ and O_2_, with dissociation constants *K*_O1_ and *K*_O2_, respectively.

Tetracycline concentrations in the cell are affected by three processes: passive diffusion across the membrane (modelled using Fick’s law), dilution due to cell growth, and active transport by TetA (modelled using Michaelis-Menten kinetics) [65]. We can combine these into the ordinary differential equation

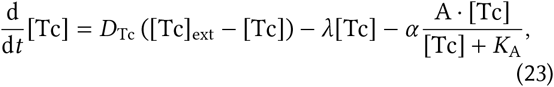

where *D*_Tc_ is the diffusion rate of tetracycline, [Tc]_ext_ is the external tetracycline concentration, *α* is the maximal transport rate per TetA molecule and *K*_A_ is the Michaelis constant. Full details of our model can be found in App. I.

We first investigate the ability of the strict and lax chemostats to capture population growth in the presence of tetracycline, applied at regular intervals of 100 and 1000 minutes (Fig. 3**b**). Comparing population and lineage estimates, we find that population growth rates are underestimated by 4–7% by single-lineage measurements, which indicates the presence of phenotypic selection. As expected, we find that the growth rate estimate increases with the chemostat size *L*, approaching an asymptotic value for *L* ≈ 50 in the strict chemostat. Due to the increased variability of the lax chemostat, it converges more slowly than its strict counterpart. We remark that a more realistic model would include (frequent) cell death, which would make single-lineage growth estimates impossible; we did not add this feature to simplify comparison.

Selection due to single-cell variability is lower when tetracycline is added more frequently, even when maintaining the same total dose over time (Fig. 3**c**). Comparing average TetA levels at different exposure frequencies (Fig. 3**d**), we see that frequent exposure causes cells to maintain high TetA levels, resulting in an effectively constitutive response that displays little cell-to-cell heterogeneity. In comparison, when tetracycline is added at large intervals, most cells have lost their TetA by the time they receive another dose, and transcriptional noise in the *tet*-operon causes higher variance in individual fitness.

Growth is inhibited most strongly at intermediate dosing intervals on the order of several cell cycles. When tetracycline is added and removed frequently, cells effectively exhibit a constitutive response that maintains low intracellular tetracycline levels throughout. In contrast, when exposure is very rare, cell growth is significantly slowed while tetracycline is applied due to the response delay, but the increased gaps between exposure windows allow population growth to rebound. Growth is lowest when exposure is rare enough to prevent a constitutive response, but still frequent enough to interrupt unrestricted growth between doses.

Fig. 3**c** shows that the single-lineage estimate significantly overestimates growth inhibition due to tetracycline at lower exposure frequencies. Following Eq. (7), the difference between the actual population growth rate and the single-lineage estimate measures the strength of selection. Comparing the behaviour of a typical forward lineage with that of a backward, or ancestral, lineage, we find that the typical ancestral lineage expresses twice as much TetA as would be observed in a mother machine (Fig. 3**d**) at low exposure frequencies. This shows that higher TetA expression is positively correlated with individual fitness when tetracycline exposure is infrequent. As we increase the dosage frequency, average TetA levels increase in all cells, which leads to less expression noise and higher TetA toxicity, and the effect disappears.

We can also probe the fitness benefit of regulation due to TetR by varying the transcription rate constant *k*_R_ (Fig. 3**e**). When *k*_R_ is large, TetR production is increased, effectively turning off the TetA-mediated response in cells. This is advantageous in the absence of tetracycline, as it prevents residual expression of the toxic efflux protein, but decreases the ability of cells to survive tetracycline exposure. In contrast, low *k*_R_ leads to “leaky” TetA expression, which hinders cell growth when tetracycline is absent, but allows cells to respond more effectively in the presence of the antibiotic. As seen in Fig. 3**e**, tuning *k*_R_ thus trades a high average growth rate in the absence of tetracycline against robust growth in stressful environments. The optimal evolutionary trade-off will depend on typical patterns of tetracycline exposure in a given environment.

We next study the effect of TetA expression variability by introducing a size parameter Ω inspired by van Kampen’s System Size Expansion [68]. Here Ω increases the TetA transcription rate *k*_A_, while simultaneously decreasing TetA export efficiency *α* and toxicity by the same amount:

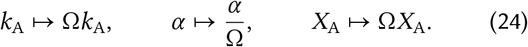

Overall, this leaves the mean (deterministic) behaviour unchanged: large Ω results in many, largely ineffective TetA molecules per cell, while small Ω results in few, but effective TetA molecules. What does change is the amount of gene expression noise, since Ω controls the copy number of TetA molecules. Small Ω causes a noisier response, which should decrease the efficacy of the operon. Fig. 3**f** confirms that small Ω does indeed result in reduced growth, but the effect only becomes noticeable below a certain threshold corresponding to a few dozen molecules per cell at maximal transcription. This suggests that moderate levels of transcriptional noise do not impose a significant fitness penalty, and may explain the noisy nature of the *tet*-response observed in nature. We remark that our model likely underestimates this fitness penalty some-what, as we do not consider delays and additional sources of gene expression noise due to separate transcription and translation.

### 3.2 Bacterial chemotaxis

We next investigate how the chemotactic strategy of a bacterial species such as *E. coli* affects its reproductive fitness. Bacteria sense their environment in order to locate nutrient sources, which they need to accumulate biomass and proliferate.

We model cells in a two-dimensional environment with periodic boundary conditions in which nutrients appear at random (Fig. 4**a**) following a homogeneous Poisson point process. Each nutrient patch diffuses with a fixed diffusion constant *D*_ν_ and decays exponentially with characteristic timescale *τ*_ν_ (Fig. 4**b**).

**Figure 4.**
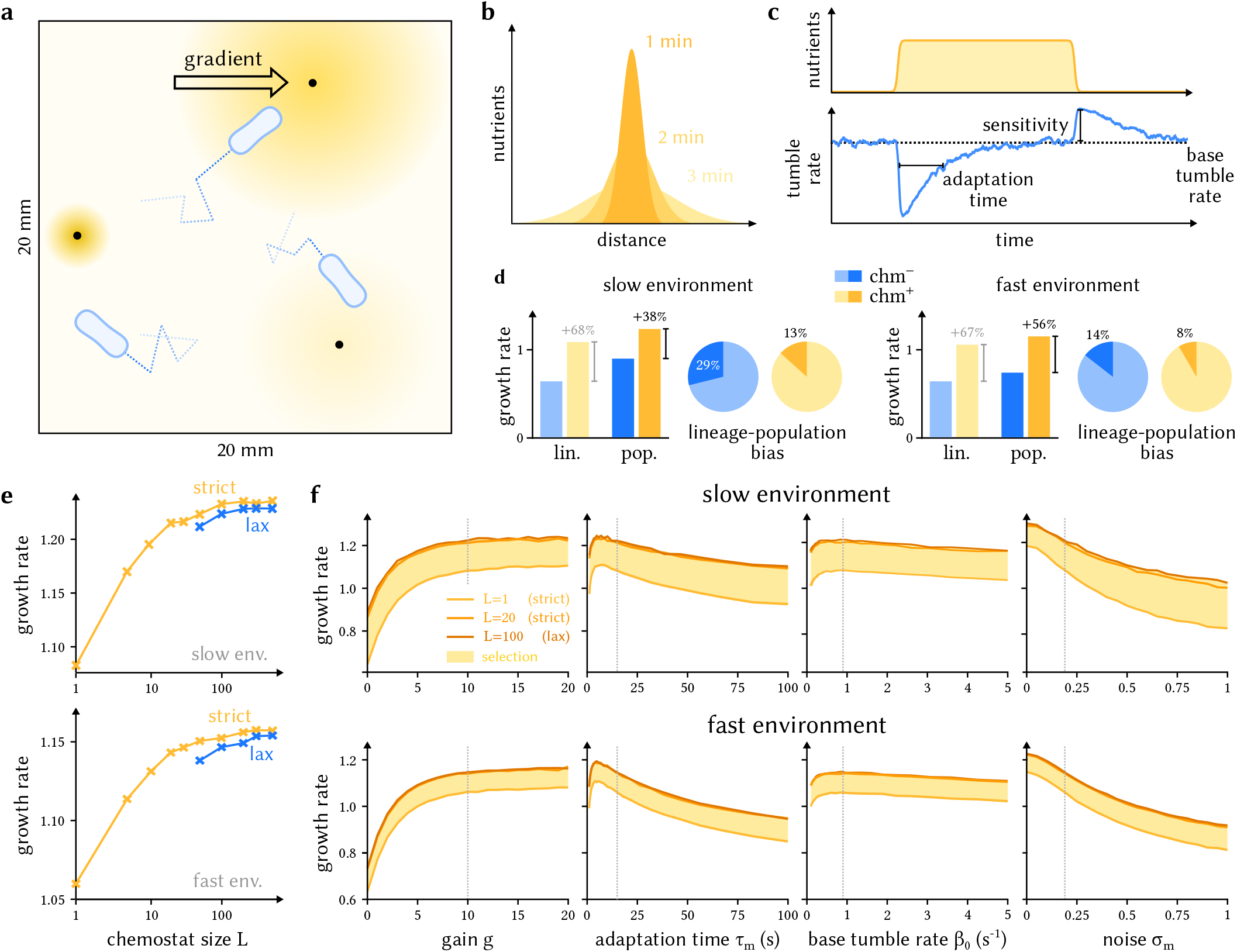
Chemotaxis in *E. coli*. **a** Illustration of a dynamic environment in which nutrient sources appear at random (not to scale). **b** Nutrients appear as point sources that diffuse and degrade over time. **c** A simple model of bacterial chemotaxis. When nutrient concentrations increase, the bacterium reduces its tumble rate, leading to longer runs in the direction of food sources. When nutrient concentrations decrease, the bacterium increases its tumble rate, reorienting more frequently to locate other sources. **d** Chemotaxis (chm^+^) increases fitness in both environments compared to a hypothetical strain with no chemotactic response (chm^−^). The difference is more pronounced in the fast environment, where nutrients degrade more quickly and directed motion becomes important for growth. Single-lineage measurements (lin.) underestimate growth rates and overestimate the benefits of chemotaxis compared to population measurements (pop.), here obtained with the lax chemostat for *L* = 100. The difference in lineage and population growth rates measures the fraction of the growth rate explained by phenotypic selection. **e** Growth rate estimates depending on the simulated population size *L* for the strict and the lax chemostat in both environments (cf. Fig. 3**b**). Estimates increase with *L*, approaching an asymptotic value for both methods. **f** Population growth rate as a function of different model parameters in both environments, for different chemostat sizes. The results for *L* = 20 and *L* = 100 are very similar, indicating convergence. The shaded area measures the selection-induced growth rate gain. Default parameters are indicated as dotted lines; optimal parameters are similar between both environments. All growth rates are given in doublings per hour. Unless otherwise noted, parameters are given in Table S2. Further model and simulation details can be found in App. J.

We implement run-and-tumble locomotion in *E. coli* following [69] using the chemotactic model in [70]. *E. coli* is a peritrichous bacterium with flagella distributed around its body. When these flagella rotate counterclockwise (CCW), they align in a bundle behind the bacterium and propel it forward, resulting in a running motion. When they rotate clockwise (CW), the bundle unravels and the cell rotates erratically, or tumbles. During runs, a cell moves with a fixed speed *v*_max_, subject to rotational diffusion with diffusion constant *D*_rot_. Tumbles occur with time-dependent rate *β*(*t*), during which the cell stops for a fixed time *τ*_tum_ and rotates randomly.

Tumbles are induced by the phosphorylated form of the protein CheY, which is rapidly dephosphorylated under normal conditions. The cell controls the tumble rate β(*t*) via the phosphorylation rate of CheY in response to external ligand concentrations (Fig. 4**c**) [71]. CheY is phosphorylated near chemotactic receptors, which are inhibited when an attracting ligand binds; this decreases the tumble rate *β* and leads to longer runs. As a consequence, the cell runs more when nutrient concentrations increase (moving up the gradient), and tumbles more when nutrient concentrations decrease (moving down the gradient), leading to reorientation. We model the tumble rate using the equation

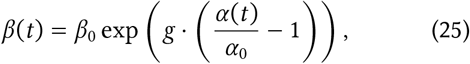

where *α*(*t*) is the current receptor activity. Here *α*_0_ is the baseline receptor activity in the absence of a signal, *β*_0_ is the corresponding baseline tumble rate and *g* is the gain or sensitivity of the system [72], see Fig. 4**c**.

The activity of a receptor depends on the external ligand concentration ν. Since the cell needs to measure gradients, or changes, in nutrient concentrations, its receptors maintain a memory of recent ligand abundances by means of methylation. Increasing nutrient levels lead to increased methylation levels *m*(*t*) at each receptor, which activates the receptor and counteracts the effect of ligand binding. For a fixed ligand concentration ν, the methylation level approaches a steady value *m*_0_(ν) with characteristic timescale *τ*_*m*_, which we model as

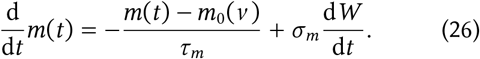

Here the second term represents white noise, which we use to model signalling noise due to fluctuations in intracellular molecule numbers [73–75]. Remarkably, the target methylation level *m*_0_(ν) depends on nutrient levels in a way to maintain the same tumble rate *β* at any constant nutrient concentration ν (robust perfect adaptation [76, 77], see below). This property allows the chemotactic system to function robustly across a large range of nutrient concentrations, reacting only to relative changes in ν [78]. The parameter *τ*_*m*_ represents the adaptation time, which determines how quickly a cell adapts to changes in nutrient concentrations, see Fig. 4**c**.

Methylation level *m*(*t*) and nutrient concentrations ν together determine the probability that each receptor is active. Since chemotactic receptors in *E. coli* such as Tar typically occur in cooperative signalling units of *h* ≈ 6, we model the activity of each cluster using Monod-Wyman-Changeux kinetics as follows:

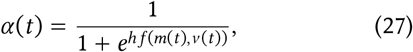

where

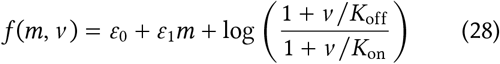

represents the free energy difference between the active and inactive state for each receptor depending on the methylation level *m* and nutrient concentrations ν. Here *ε*_0_ and *ε*_1_ represent the effect of methylation, and *K*_on_ and *K*_off_ are the ligand dissociation constants for the active and inactive state of the receptor, respectively. Requiring that the receptor activity *α*(*t*) is equal to the baseline *α*_0_ in the absence of concentration changes (robust perfect adaptation) determines the steady-state methylation level *m*_0_(ν) implicitly via

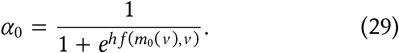

To model growth, we assume that cells accumulate biomass at a rate depending on the current nutrient concentration ν following the Monod equation

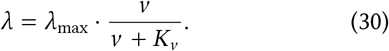

Here λ_max_ is the maximal growth rate of a cell, and *K*_ν_ is the nutrient concentration necessary for half-maximal growth. Division occurs once a cell reaches a fixed volume *V*_*d*_. Cells will therefore grow faster when they are located in regions of high nutrient concentration ν. Full details of the model can be found in App. J.

We first quantify the fitness benefit due to chemotaxis in comparison with a non-chemotactic mutant in which the signal gain is zero. These modified cells still perform run-and-tumble motion, but do not incorporate any information about local nutrient concentrations. We estimate the long-term growth rates in two different environments (“fast” and “slow”), which differ in the frequency at which nutrients appear and are degraded (Fig. 4**d**). Sensing increases population growth rates by 38% in the slow environment, and 56% in the fast environment. While the exact numbers biologically depend on the cell strain and environment, this supports the view that chemotaxis substantially increases a population’s nutrient intake and proliferation rate in a fluctuating environment.

As with the tetracycline model, single-lineage tracking substantially underestimates population growth rates for both strains in both environments (Fig. 4**d**). As cells move around the environment, the random nature of their run-and-tumble motion and the location at which nutrient patches appear mean that some cells acquire many more nutrients than others, leading to phenotypic selection even in otherwise identical cells. This phenomenon is not visible when tracking a single lineage and yields biased growth estimates in this case. Phenotypic selection acts as a buffer that decreases the fitness benefit associated with chemotaxis, as even non-chemotactic cells can occasionally end up in nutrient-dense regions and proliferate rapidly for some time. Thus even in the absence of sensing, having cells spread throughout the environment provides a systematic fitness advantage, a phenomenon reminiscent of bet-hedging. As a result, single-lineage measurements overstate the growth rate gain due to chemotaxis (Fig. 4**d**). We quantify selective effects by comparing growth rate estimates for chemotactic bacteria in chemostats of different sizes (Fig. 4**e**). As predicted, and as observed in the tetracycline model, the estimates increase with *L* and appear to converge to a limiting value.

We next investigate how chemotactic parameters affect fitness in both environments (Fig. 4**f**). The gain *g* amplifies the signal received from the cell’s receptors, allowing the cell to react more strongly to local nutrient gradients, at the cost of also amplifying the noise in the circuit. As for TetA copy number in Fig. 3**f**, increasing *g* generally improves fitness, but the difference is small beyond *g* ≈ 10, corresponding to the apparent Hill coefficient reported in [72]. In an idealised chemotactic circuit with no noise, higher gains led to a more marked improvement (not shown).

The adaptation time *τ*_*m*_ controls the timescale over which the cell measures gradients. A small value of *τ*_*m*_ allows the cell to respond quickly to changes in nutrient levels, but limits the accuracy with which the cell can measure gradients; conversely, a larger *τ*_*m*_ delays the cell’s response, but allows it to estimate gradients with higher precision. Fig. 4**f** suggests that an intermediate adaptation time of several seconds is optimal in both environments. This is somewhat below the value *τ*_*m*_ ≈ 15 s reported in [79], noting that response times vary significantly between cells [75]. The optimal adaptation time likely depends on mechanistic parameters like movement speed, tumble duration and rotational diffusion, as well as the nature of rotations in two vs. three dimensions.

The baseline tumble rate *β*_0_ had a comparatively small effect on chemotactic performance (Fig. 4**f**). When the chemotactic signal is strong, changes in local nutrient concentrations are amplified by a factor of *g* ≈ 10 to modulate the tumble rate, which largely determines movement; conversely, when the cell is not moving in a gradient and the chemotactic signal is weak, increasing or decreasing the tumble rate does not make a big difference.

In our model, the strength of methylation noise *σ*_*m*_ strongly influences chemotactic performance (Fig. 4**f**). This is not surprising, as noise impedes cells’ ability to reliably convert a signal into locomotion [80]. In the slow environment in particular, increased noise furthermore leads to increased selection, as it increases the variability in behaviour between cells. Similar to the non-chemotactic variant in Fig. 4**d**, this slightly buffers the fitness penalty due to decreased sensing.

## 4 Discussion

We present two related Monte Carlo algorithms to analyse population behaviour in agent-based models of bacteria, the strict and the lax chemostat. Both algorithms achieve quadratic computational complexity by randomly tracking lineages in a population and can be seen as particle filters, a form of Sequential Monte Carlo [28, 29]. The strict chemostat resembles a generalised Moran process and is closely related to cloning algorithms in statistical physics [30, 31]. Restricting the population size introduces a finite-size bias into our estimates, which is most pronounced when extrapolating from a single lineage. This bias is of order 1/*L* in the simulated population size *L*, and can be controlled by increasing the size of the chemostat. While increasing only the simulation time *t* or only the population size *L* does not result in statistically consistent estimates, increasing both in tandem does.

Our results are based on the approach to microbial population dynamics presented e.g. in [36, 81], which expresses the population growth rate Λ in terms of the statistical behaviour of individual lineages. This connection also underlies the original cloning algorithms [30, 31], and can be used to quantify the bias in estimating growth rates from a restricted population, e.g. a single lineage [27]. We show that this phenomenon stems from the inability of single lineages to capture selection within a population, leading to consistent underestimation of growth rates. By simulating multiple competing lineages in a restricted population, we can capture the growth-rate-increasing effects of selection in the long run.

We illustrate these algorithms by applying them to two biophysical models of bacterial behaviour: the stochastic response of the *tet*-operon, an adaptive antibiotic resistance mechanism in *E. coli*, and chemotactic foraging of bacteria in a random environment. In both cases, single-cell variability due to molecular noise results in heterogeneous outcomes for individual cells, leading to measurable selection effects between otherwise identical cells. Our approach allows us to predict how biophysical parameters such as transcription rates in the *tet*-operon, or sensitivity and tumble bias in chemotaxis, affect population fitness, and to quantify the role of variability and phenotypic selection within populations [25, 37].

The algorithms presented in this paper are subject to several limitations. Bacterial growth is rarely exponential in the long run, due to effects including crowding and resource depletion. Nevertheless, exponential growth rates can predict behaviour in size-restricted populations [6, 38], and are commonly used measures of bacterial fitness [7, 8, 13–15]. More critically, we neglect direct cell-to-cell interactions such as quorum sensing that can play a large role in determining population behaviour; extending our results to include these effects will be of significant interest for realistic single-cell modelling.

Nonetheless, our results suggest that population behaviour can be extrapolated from surprisingly few cells at once, on the order of 20–100 in our applications; this is orders of magnitude fewer than typical bacterial populations, and within the realm of computational tractability. While we expect the exact numbers to depend on the application, it is likely that more sophisticated statistical approaches [82] will enable further improvements in predicting bacterial population behaviour from mechanistic single-cell models.

## Acknowledgements

This research was funded by an Andrew Sisson Grant to KÖ and an ARC Laureate Fellowship to MPHS (FL220100005), and supported by the University of Melbourne’s Research Computing Services and the Petascale Campus Initiative. The authors would like to thank Daniel Schultz, Augustinas Sukys, James Holehouse, Irina Kalita and Douglas Brumley for valuable discussions.

## Code availability

The algorithms presented in this paper are available as the Julia package Chemostats.jl at

https://github.com/kaandocal/Chemostats.jl

The models used for numerical evaluation can be found at

https://github.com/kaandocal/chemostat_models

## Appendix

### A Branching processes

There are many ways to mathematically describe population growth, generally falling under the umbrella of branching processes. For our derivations, we will restrict ourselves to Markovian multitype branching processes, which illustrate the essential ideas without becoming mathematically unwieldy. Our applications are more complex, featuring age-dependent growth in a fluctuating environment, and the proofs that follow can be adapted to this more general scenario. For a general overview of branching processes we refer to [83].

In a multitype branching process, we assume that there are finitely many possible types *i* = 1, …, *s* of individuals. Mathematically, the type of each individual cell can include its birth size and growth rate [39, 43], ribosome and protein abundances [21, 84], spatial position [85] or responsiveness to antibiotics [47]. Note that this notion of cell type is more general than the biological definition of a cell type.

Each cell of type *i* has an exponentially distributed lifetime with rate λ_*i*_, at the end of which it either dies with probability *d*_*i*_, or divides into two daughter cells of type *j* and *k* with probability *b*_*jk,i*_. Here the order of the two daughter cells *j* and *k* does not matter, so *b*_*jk,i*_ = *b*_*kj,i*_. The probabilities for all these events (death and division) sum up to one, so that

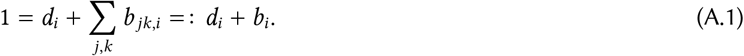

Since every cell has an exponentially distributed lifetime, this population process is memoryless (Markovian): the future of the population only depends on the number of cells *N*_*i*_(*t*) present at a given time *t*. The expected number *N* _*i*_ of individuals of type *i* is described by the evolution equation

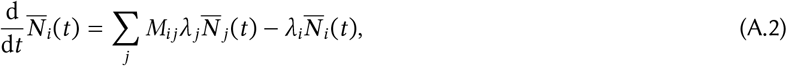

where

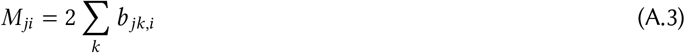

is the next-generation matrix. Here the first term in Eq. (A.2) represents newly born cells of type *i*, and the second term represents cells of type *i* that die or divide. Since Eq. (A.2) is a linear system of ODEs, we can expand the solution in terms of eigenvectors:

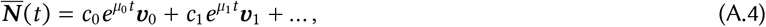

where Re*μ*_0_ ≥ Re*μ*_1_ ≥ … are the eigenvalues sorted by decreasing real part, the *v*_*k*_ are the corresponding eigenvectors, and the *c*_*k*_ are coefficients that depend on the initial condition 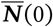. Under standard assumptions such as irreducibility, one can show that the largest eigenvalue *μ*_0_ is real and equal to the population growth rate Λ; we will assume that Λ > 0, so that we are dealing with a growing population. Computing the growth rate Λ therefore reduces to computing the maximal eigenvalue of a linear operator. For more complex (age-dependent) branching processes, the growth rate Λ is determined implicitly by a matrix Euler-Lotka equation.

We will also need the cell type distribution

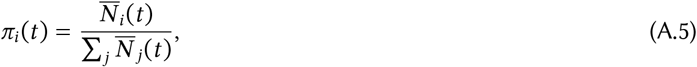

which satisfies

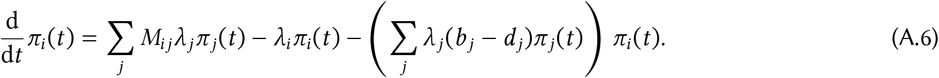

Here the difference to Eq. (A.2) is in the additional last term, which ensures that the probabilities sum to 1 at all times. For large *t*, we can use the expansion Eq. (A.4) to show that

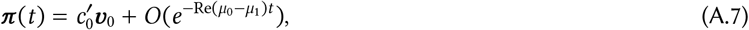

where 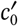 is a constant and Re*μ*_0_ −Re*μ*_1_ is the spectral gap between the leading two eigenvalues. Therefore, if we normalize *v*_0_ so that it sums to 1, we see that *π* := *v*_0_ represents the steady-state distribution of cell types in the population, and the distribution π(*t*) converges to this steady state exponentially quickly. For later reference, we observe that the eigenvalue property of *π* implies

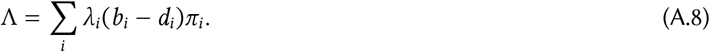

The above discussion yields the expected (large-scale) behaviour of a population, ignoring stochastic fluctuations. Under very general conditions, assuming Λ > 0, the behaviour of any individual realisation is dictated by the Kesten-Stigum theorem (see e.g. [86]), which states that

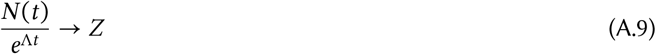

converges to a finite number *Z* almost surely. Thus, with probability 1 the observed population size grows exponentially for large *t*, with growth rate predicted by the deterministic equations. We note that *Z* can be 0 for some runs, corresponding to the population going extinct.

Taking expectations and comparing with Eq. (A.4) shows that E[*Z*] = *c*_0_, since we assume that *v*_0_ sums to 1. If we start with one individual of type *i*, E[*Z*] is the reproductive value of type *i* [83]. When the initial individual is sampled from the steady-state distribution *π*, one can show that E[*Z*] = 1, but in general, E[*Z*] is no easier to compute than Λ.

### B Averaging growth rate estimates

In Section 2.1 we considered the effect of averaging independent growth rate estimates 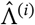 to reduce sampling error. Taking the arithmetic mean of the growth rates is equivalent to taking the geometric mean of the population sizes:

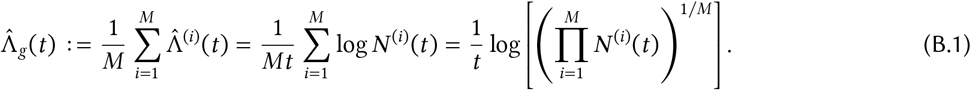

We therefore call this the geometric mean estimator. Here we assume that each population starts with one individual. One sees from Eq. (4) that this estimator is biased since E[log *Z*] ≠ 0 in general. In the simple case of the linear birth process, we show in App. G that E[log *Z*] ≈ −0.577.

Another approach briefly discussed e.g. in [87] uses the arithmetic mean of the population sizes, resulting in

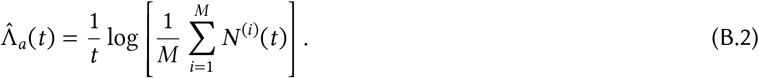

We call this the arithmetic mean estimator; the AM-GM inequality implies that 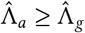. Fixing *t*, in the limit of large *M* the arithmetic mean estimator is approximately

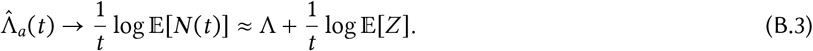

As follows from our discussion in App. A, E[*Z*] ≠ 1 in general, so like the geometric mean, the arithmetic mean estimator remains asymptotically biased as *M* → ∞, with bias of order *O*(1/*t*). For the special case of the linear birth process, E[*Z*] = 1 and this estimator is asymptotically unbiased, although it remains biased for finite *M*.

We remark that Eq. (B.2) simply estimates the growth rate of the population obtained by pooling the *M* populations, so it is equivalent to starting with a larger initial population. From our discussion in App. A, E[*Z*] = 1 is a more natural condition than E[log *Z*] = 0; the former occurs if the initial individual is drawn from the steady-state distribution over cell types. Furthermore, the geometric mean estimator yields −∞ if any population goes extinct (in which case E[log *Z*] = −∞). For these reasons we generally recommend the arithmetic mean estimator in practice.

### C Lineage statistics

In this section we discuss the relation between lineage and population statistics following [25]. Consider a microbial population tree originating from a single individual (Fig. S1**a**). Every cell at a fixed time *t* has an ancestral lineage, and these ancestral lineages together make up the population tree.

To interpret Eq. (5), we define two probability distributions over these lineages. The forward distribution starts at the original ancestor and follows its descendants in time. Each time our cell divides we pick a daughter cell at random, until we end up at a cell that has not yet divided (for simplicity, we ignore cell death here). At every division event, we have a probability 1/2 of sampling either, so the probability of sampling a lineage *L* with *n*(*t*) divisions is simply

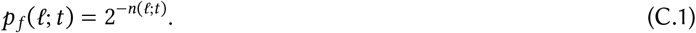

We remark that the forward distribution is the one obtained in a mother machine, where a random daughter cell is discarded at each division. Multiplying by 2^*n*(*t*)^ and summing over all *N* (*t*) lineages *L*_1_, …, *L*_*N* (*t*)_ yields

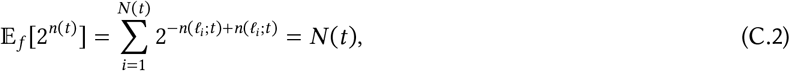

which shows Eq. (5). In particular, we can interpret 2^*n*(*L*;*t*)^ as the absolute fitness of a lineage, as measured by its contribution to the total population.

The forward distribution assigns different probabilities to each individual in the current population, and thus does not represent the statistics of a randomly sampled individual. For this reason, we define the backward distribution over lineages by picking the ancestral lineage of an individual at time *t* at random. Since each ancestral lineage has the same probability 1/*N* (*t*), we obtain

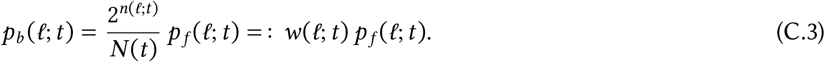

The term *w*(*L*; *t*) represents the relative fitness of each lineage [25]. Lineages that have divided more often than others are fitter, and contribute relatively more offspring to the population. Comparing Eq. (C.1) and Eq. (C.3) yields

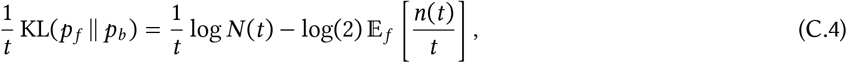

and we obtain Eq. (7) by taking *t* → ∞. For this reason, we view the difference between the backward and forward distributions as measuring the amount of selection in a population; for more details, we refer to [25, 37, 81].

### D Analysis of the lax chemostat

In this section we show that Eq. (10) yields unbiased estimates of the true population size *N* (*t*). This is a common property of particle filters, of which the lax and the strict chemostats are examples [28, 29]. The key property is that the death rate *δ*(*t*) on the interval (*t*_*k*_, *t*_*k*+1_) only depends on the behaviour of the population before time *t*_*k*_. Based on this observation, we will show by induction on *k* that Eq. (10) is unbiased.

Starting with an initial death rate *δ*_0_ ≥ 0 on (0, *t*_1_), assume that the unthinned population consists of the individuals *x*_1_, …, *x*_*n*_ at time *t*_1_. For times *t* > *t*_1_, the total population is made up of the offspring of these *n* lineages, so we can write

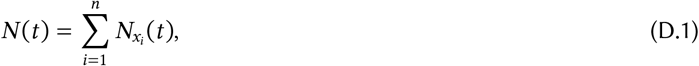

where 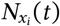 is the size of the population descended from cell *x*_*i*_. The probability that the lineage leading up to cell *x*_*i*_ has not been culled by time 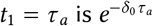, so we can rewrite Eq. (D.1) as

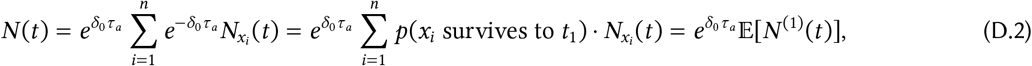

where *N* ^(1)^(*t*) is the size of the population descended from the individuals *x*_*i*_ that survive until time *t*_1_. We roughly illustrate the idea behind this procedure in Fig. S1**b** for the related case of the strict chemostat.

**Figure S1.**
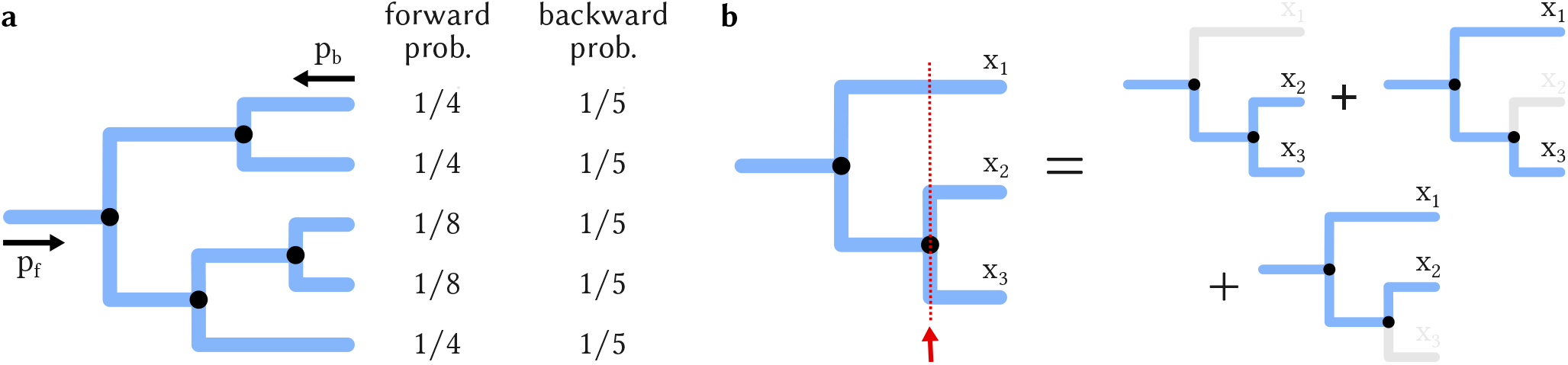
Relating population and lineage statistics. **a** Forward and backward distribution in an example population tree. Starting from the original ancestor, the forward probability of a lineage halves with each division as we track either daughter with 50% probability. Cells in the final population can have different forward probabilities depending on the number of ancestors they have. In contrast, the backward distribution treats each cell in the population equally. **b** Decomposing a population as a statistical average of subpopulations. When the cell count reaches 3 (dotted line), we randomly discard a cell and track the remaining subpopulation. Since each cell has probability 2/3 of being kept, the subpopulation will on average be 2/3rds the size of the full population. This can be repeated at each cell division to extrapolate *N* (*t*) from a limited number of lineages. When restricted to a single cell at a time, this argument yields Eq. (5).

We can repeat this argument for the next interval (*t*_1_, *t*_2_), replacing the full population by the subpopulation descended from the cells still present at time *t*_1_. Proceeding the same way, we obtain

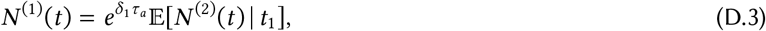

where *N* ^(2)^(*t*) is the size of the population descended from the lineages that survive until time *t*_2_. Here we take expectations conditional on the chemostat at time *t*_1_, since *δ*_1_ depends on the observed population at time *t*_1_ via Eq. (15). As a result, for *t* > *t*_1_ we have

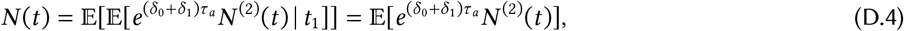

using the law of total expectation. We can repeat this for subsequent intervals: the same argument as before yields the general relation

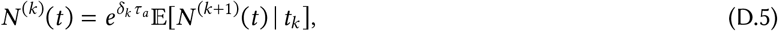

for any *k* ≥ 1, and for *t* > *t*_*k*_ we get

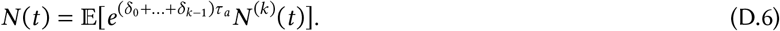

Assuming the observation time *t* is in the *k*-th interval (*t*_*k*_, *t*_*k*+1_), where we use the thinning rate *δ*_*k*_, the observed population size on average equals

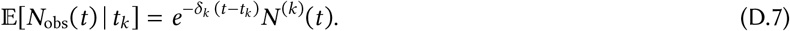

Combining this with the previous equation we obtain

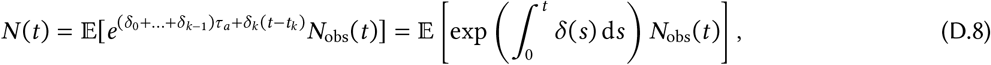

which establishes the unbiasedness of Eq. (10).

Our proof works for any piecewise constant function *δ*(*t*) that is estimated from the history of the chemostat. The formula Eq. (15) uses the estimated growth rate 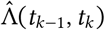 on the previous interval, which can be unreliable if the interval is short. For more stable estimates, we can equally well use moving averages of the form

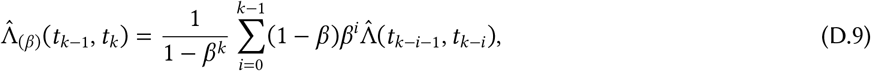

where 0 ≤ *β* < 1 represents a tuning parameter. Here for *β* = 0 we recover Eq. (13), whereas *β* > 0 takes into account the empirical growth rate at earlier times.

The lax chemostat therefore yields an unbiased estimate of *N* (*t*). As discussed in the main text, Jensen’s inequality implies that it will systematically underestimate the growth rate compared to tracking the full population:

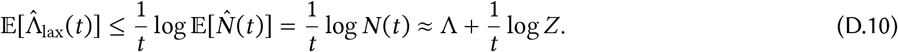

This nonlinearity bias is caused by fluctuations in estimating *N* (*t*) from 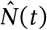 [27]. We can estimate this error by writing

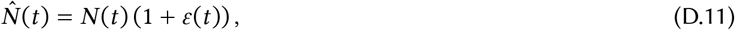

where *ε*(*t*) represents these fluctuations, which average out to 0. Assuming that *ε*(*t*) is small, we can Taylor expand the logarithm to find

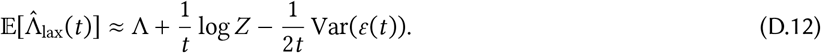

Here the last term represents the additional bias incurred by subsampling the population. Unfortunately, the variance of *ε*(*t*) grows with *t*, so this term does not disappear in the limit *t* → ∞: the lax chemostat remains biased for long times.

Deriving accurate estimates for this bias requires analysing the statistical behaviour of the lax chemostat, which is not straightforward. For large *L*, however, the lax chemostat maintains a more or less constant population size, and behaves similarly to the strict chemostat. In the next section, we will study the strict chemostat and show that the introduced long-term bias is on the order of 1/*L*. We therefore expect the lax chemostat to exhibit the same systematic bias of order 1/*L* for large *L*. This is borne out by our numerical benchmarks (see Fig. 3 and Fig. 4). Since the observed population size regularly dips below *L*, we note that the lax chemostat will incur a slightly larger estimation error than its strict counterpart for fixed *L*.

Similarly, analysing the variance of the lax chemostat estimator we find

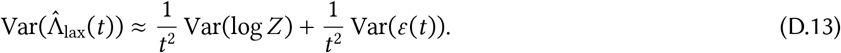

The fact that the long-term bias in Eq. (D.12) is on the order of 1/*L* suggests that

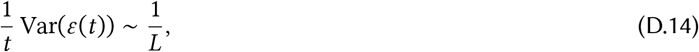

which in turn yields

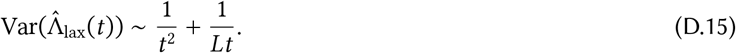

From this we can derive Eq. (12) using the bias-variance decomposition

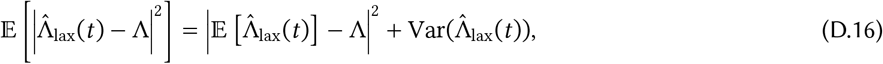

and using the fact that the bias is of order *O*(1/*t* + 1/*L*).

### E Analysis of the strict chemostat

#### E.1 Unbiased sampling

Much like the lax chemostat, the strict chemostat yields unbiased estimates of the full population size *N* (*t*) (more precisely, in the presence of cell death, it yields an unbiased estimate of E[*N* (*t*)]). Our proof in this section can be seen as an extension of Eq. (5), which corresponds to the case *L* = 1. We first consider the case without cell death, which follows essentially the same reasoning as in the previous section. Eq. (16) clearly holds until the first time of division, since *B*(*t*) = 0 and the full population size equals *L*. If we denote the first division time by *t*_1_, we have *L* + 1 lineages *x*_1_, …, *x*_*L*+1_, and Eq. (D.1) from the previous section holds with *n* = *L* + 1. The probability that a lineage will be kept in the chemostat is now *L*/(*L* + 1), so using the same argument as for the lax chemostat we can write

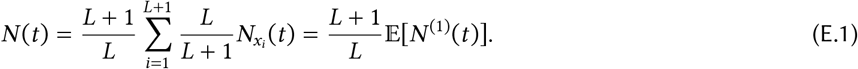

We illustrate the idea behind this argument in Fig. S1**b**. As we did before, we can iterate this procedure to obtain

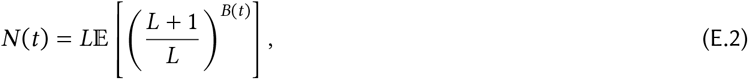

on the interval (*t*_*k*_, *t*_*k*+1_) where *B*(*t*) = *k*. This shows that Eq. (16) is unbiased in the absence of cell death.

The case where cells can die is slightly more complicated; here we must assume *L* > 1. The strict chemostat replaces a dead cell by an independent copy of one of the remaining *L* − 1 cells at random, which adds cells to the population. Assume for illustration that the first event is a death at time *t*_1_, with surviving lineages *x*_1_, …, *x*_*L*−1_. Then Eq. (D.1) holds with *n* = *L* − 1. Let *x*_*L*_ be a copy of one of the other lineages, so we can write

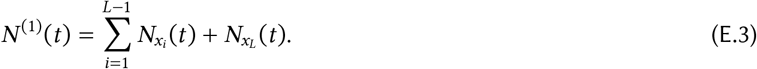

Here the cloned lineage *x*_*L*_ produces its own descendants independently of the others. Since each lineage has probability 1/(*L* − 1) of being cloned, we have

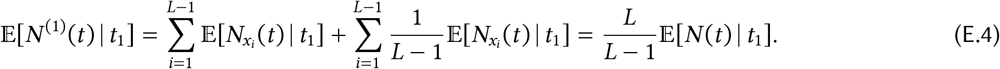

Here we use 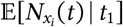, the *expected* population size at time *t* descended from lineage *x*_*i*_. With this small modification we can proceed as above, accounting for division and death events as they occur, and considering the *expected* population sizes E[*N* (*t*)] and E[*N* ^(*k*)^(*t*)]. This shows that Eq. (16) provides an unbiased estimate of E[*N* (*t*)], the *expected* population size at time *t*.

We can modify the strict chemostat to avoid cloning cells. In this case, the observed population size *N*_obs_(*t*) can drop below *L* when a cell dies. When this happens, we allow the population to recover, only removing cells when the population size exceeds *L*. Adapting the above argument to this case, we derive the estimate

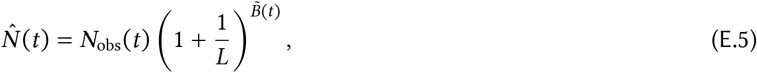

where 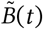 only counts those birth events where a cell is removed from the population, i.e. only birth events occurring when the population size is already *L*. We provide an implementation of the strict chemostat with and without this modification in App. F.

#### E.2 Error estimates

The strict chemostat is closely related to cloning algorithms [30, 31] and population control Monte Carlo, for which asymptotic error estimates are available [32, 88]. In this section, we rederive these for models of cell growth, referring to e.g. [89] for rigorous error bounds. In keeping with App. A, we assume that our population is a Markovian multitype branching process to simplify our argument. In this scenario, the strict chemostat is a Markov chain, with state represented by the number of cells *n* = (*n*_1_, …, *n*_*s*_) of each type.

If there are *n*_*i*_ cells of type *i*, the total rate at which a cell of type *i* dies is *n*_*i*_λ_*i*_*d*_*i*_. Whenever this happens, it is replaced by a random copy of one of the remaining *L* − 1 cells. The copied cell is of the same type *i* with probability (*n*_*i*_ − 1)/(*L* − 1), and of a different type *j* ≠ *i* with probability *n*_*j*_ /(*L* − 1). Similarly, the total rate at which a cell of type *i* divides into cells of type *j* and *k* equals *n*_*i*_λ_*i*_*b*_*jk,i*_. When this happens, the dividing cell is replaced by its daughter cells, and a random one of the *L* + 1 remaining cells is removed, say of type *l*. Considering all possibilities for *i, j, k* and *l*, we see that the strict chemostat is described by the following master equation:

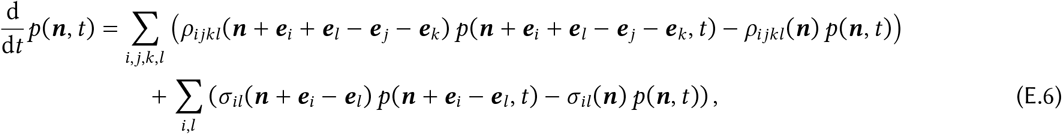

with propensities

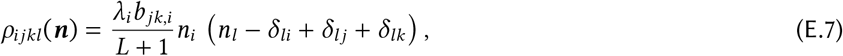

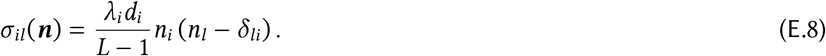

Here *p*(*n, t*) is the probability that the chemostat is in state *n* at time *t*, and *e*_*i*_ is the *i*-th unit vector (zero everywhere with 1 in the *i*-th position).

Following [32] we analyse the behaviour of the strict chemostat in the limit of large *L* using van Kampen’s System Size Expansion [68, 90, 91]. Here we express the cell count *n*_*i*_ as a sum of a deterministic part *x*_*i*_ and a noise term *ξ*_*i*_:

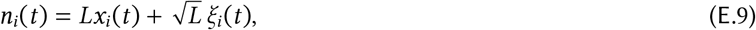

with *x*_*i*_ and *ξ*_*i*_ of order *O*(1) in the system size. The total abundance of cells scales with *L*, whereas the size of fluctuations scales with 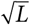. Since the total number of cells equals *L*, we require

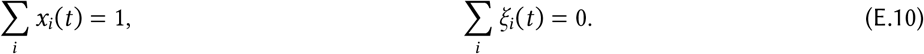

Performing the system size expansion as outlined e.g. in [32, 91], we find that the deterministic variables *x*_*i*_(*t*) evolve according to Eq. (A.6). Thus in the limit of large *L*, the distribution of cell types in the chemostat follows that of an unrestricted full population, *x*_*i*_(*t*) = *π*_*i*_(*t*), with fluctuations represented by *ξ*. For the noise variables *ξ*_*i*_, one can derive the linear SDE

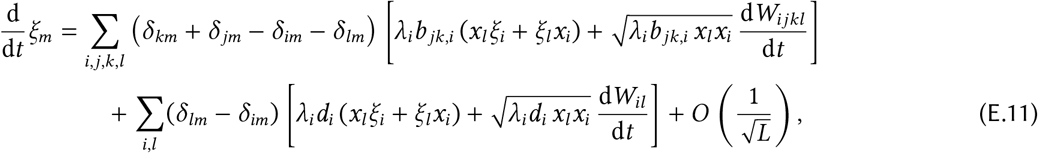

where the *W*_*ijkl*_ and *W*_*il*_ are independent Wiener processes. As a result, for deterministic initial conditions *ξ* (0) = 0 we find that

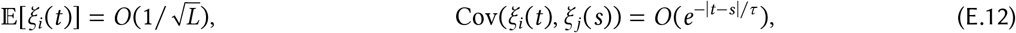

where *τ* = (Re*μ*_0_ − Re*μ*_1_)^−1^ is the relaxation timescale of the macroscopic population (App. A). Altogether we obtain

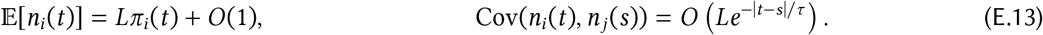

The corrections to the mean are of order *O*(1), see e.g. [92]. As a consequence, we can compute population averages with error of order *O*(1/*L*) in a chemostat, without simulating an exponentially growing population.

We can now derive error estimates for the strict growth rate estimator Eq. (17), which can be written as

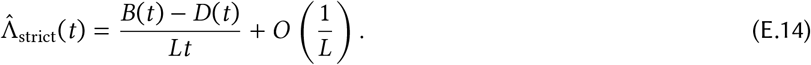

It suffices to show that

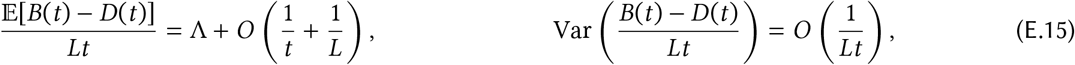

for large *L* and *t*. This implies that the bias in Eq. (17) is asymptotically of order *O*(1/*t* + 1/*L*), and the variance of order *O*(1/*Lt*), which yields the asymptotic error estimates presented in the main text.

To do so, we decompose all birth and death events into individual reaction channels:

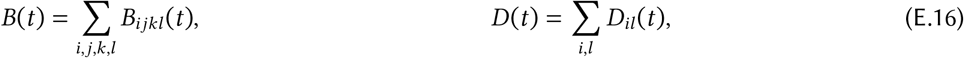

with propensities ρ_*ijkl*_ and *σ*_*il*_ defined in Eqs. (E.7) and (E.8). The Poisson time-change representation [93] of the master equation Eq. (E.6) implies that the compensated counting processes

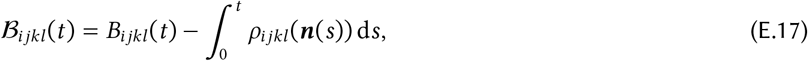

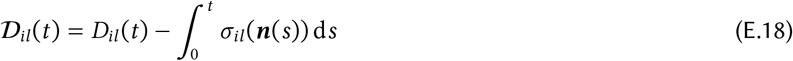

are martingales. In particular,

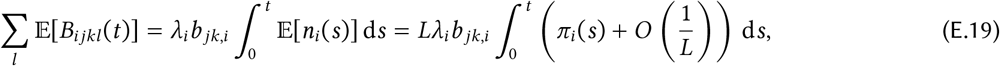

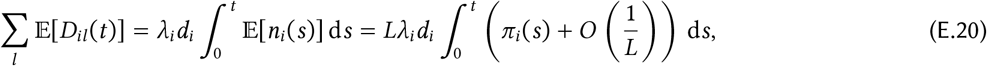

where we used Eq. (E.13). Summing over *i, j, k* and comparing the time-dependent solution *π*_*i*_(*t*) with its steady-state limit *π*_*i*_, we can write

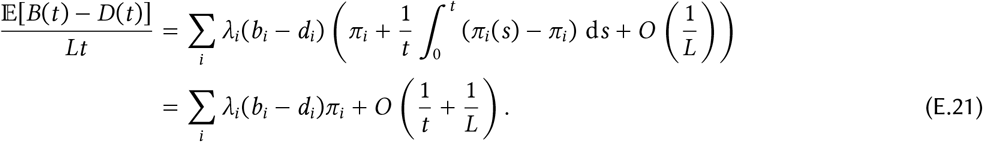

Here we use the fact that *π*_*i*_(*t*) converges to *π*_*i*_ exponentially quickly. The identity Eq. (A.8) then yields

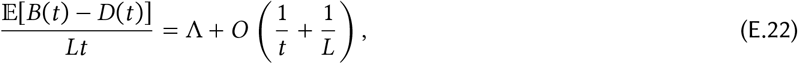

which shows the first part of Eq. (E.15).

For the variance estimate, we note that the quadratic variation of the compensated counting processes Eqs. (E.17) and (E.18) yields

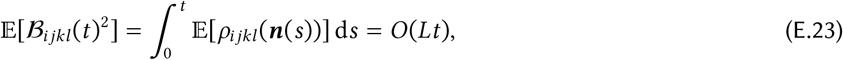

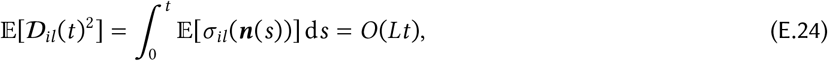

and since two reactions cannot occur simultaneously, the compensated processes are uncorrelated [93, 94], so that

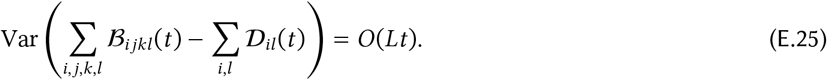

Furthermore, using the Linear Noise Approximation Eq. (E.9) together with Eq. (E.13) we can estimate

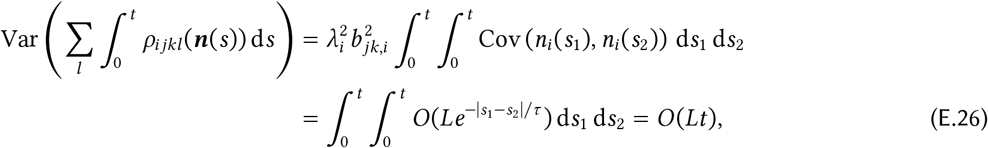

and similarly for all (co)variances including birth and death reactions. Here *τ* is the relaxation timescale of the full population, see Eq. (E.13). As a result, by the Cauchy-Schwarz inequality we obtain

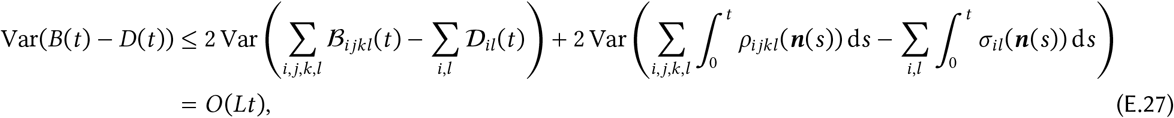

which implies the second part of Eq. (E.15).

### F Implementation

This section contains implementation notes for the algorithms used in this paper. Pseudocode for the lax and strict chemostats is provided in Algorithm 1 and Algorithm 2, and a Julia implementation can be found at https://github.com/kaandocal/Chemostats.jl.

#### Direct simulation

To simulate a full population until time *T*, we maintain one global queue *Q* of “live” cells. At each step, we remove one cell from this queue and simulate it until we reach the end of its lifetime, or the maximal simulation time *T*. If the cell divides, both daughters are placed in the queue to be simulated later; if it reaches the maximal simulation time, the cell is saved as belonging to the population at time *T*. We iterate this procedure until the queue is empty. The population size *N* (*T*) is obtained by counting the number of cells saved. We remark that the order of simulation does not necessarily match physical time, which does not affect the results in our scenario (as we assume that cells do not directly interact). In particular, we can simulate multiple cells from the queue in parallel.

#### Thinning

Here we proceed as above, but apply an exponential death rate *δ* to each cell. That is, if a cell is simulated until age *a*, we keep it with probability *e*^−*δa*^ and discard it otherwise.

#### Lax chemostat

We fix a target size *L*, an adaptation time *τ*_*a*_ and optionally a smoothing parameter 0 ≤ *β* < 1. We simulate a thinned population with constant death rate on each interval (*t*_*k*_, *t*_*k*+1_), starting with *δ*_0_ = 0 in the first interval. At the end of each interval, we estimate the current growth rate either using Eq. (13), or the running average Eq. (D.9). We then compute the next value of *δ*_*k*+1_ using Eq. (15), and continue simulating the population with this new thinning rate. Note that we can simulate cells in parallel on each time interval (*t*_*k*_, *t*_*k*+1_), only synchronising at the update times to compute 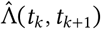.

#### Strict chemostat

This requires more care. Since we discard (resp. clone) a random cell whenever any cell in the population divides (resp. dies), we must keep cells synchronised at all times (this is what makes parallelisation difficult). One approach is to simulate all cells in tandem, as done e.g. in [95], but that mixes single-cell and population handling, which may be undesirable. We take a different approach, maintaining a priority queue *Q* of cells sorted by the current simulated time *t*(*c*) of each cell *c*. At each step, we take the earliest cell and simulate it until we reach the end of its lifetime, or the maximal simulation time *T*, after which we *reinsert* the cell into the priority queue (with its current time). When we sample a cell at time *t* that has already been simulated, this guarantees that all other cells have been simulated past time *t*. Furthermore, all cells in the queue were alive at time *t*, since we reinsert cells into the queue before removing them. If the current cell divides, we place both daughters in the priority queue and end up with *L* + 1 cells alive at time *t*, of which we discard one at random before proceeding. If the sampled cell dies without leaving offspring, we instead copy one of the remaining *L* − 1 cells at time *t* and place it in the queue. The cloned cell is simulated independently starting at time *t*; implementing this requires saving the history of each cell. Once we reach time *T*, all cells in the remaining queue belong to the population at time *T*.

For the single-lineage estimator, this approach simplifies as we only need to keep track of one cell at any given time. Whenever the simulated cell divides in two, we discard one of the daughter cells and continue simulating the remaining daughter, until we either reach time *T* or the current cell dies without leaving offspring.

##### Algorithm 1

Implementation of the lax chemostat.

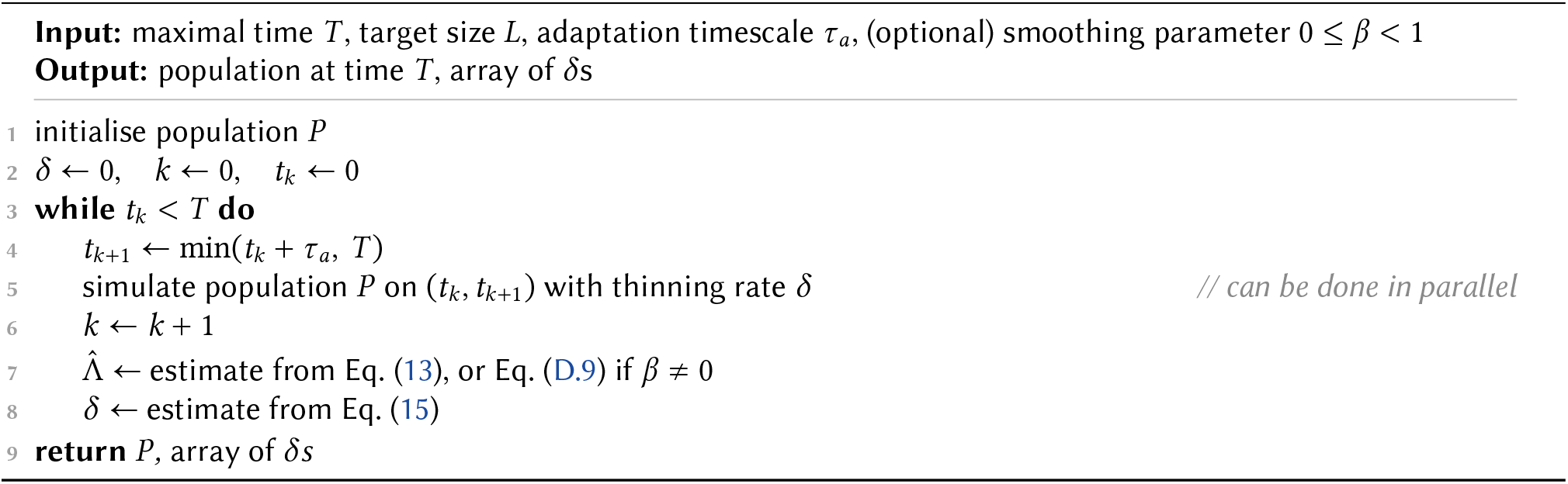

##### Algorithm 2

Implementation of the strict chemostat.

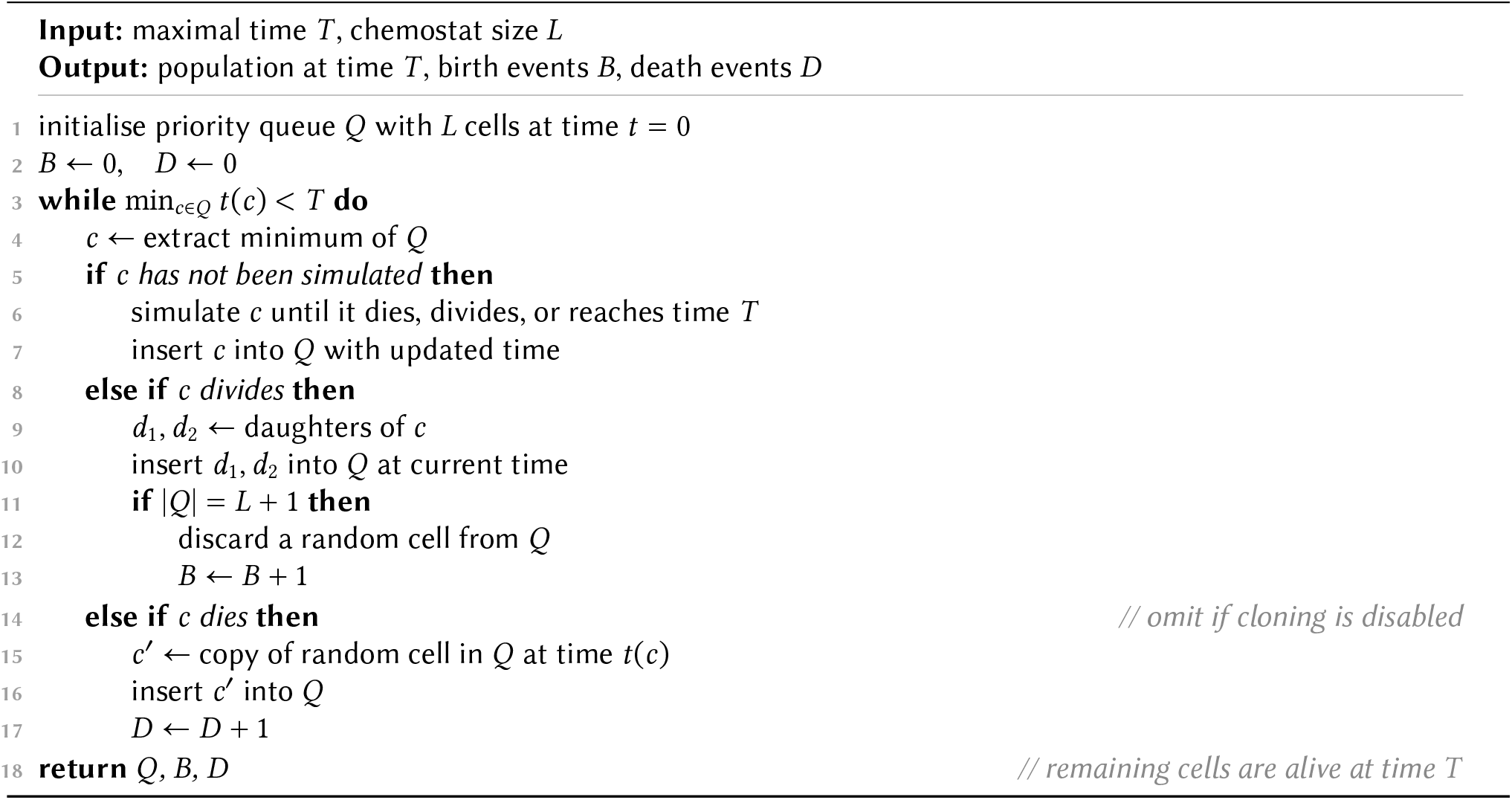

### G Linear birth process

The linear birth process, also known as the Yule process, describes a population of cells whose lifetimes are independent and exponentially distributed with rate Λ (here the population growth rate equals the single-cell division rate). Due to its simplicity, we can use the linear birth process as an analytically tractable illustration of some of the concepts discussed in this paper.

Since every cell in this process is memoryless, the state of the population is entirely determined by the number of individuals *N* (*t*). The evolution of this random quantity is described by the master equation

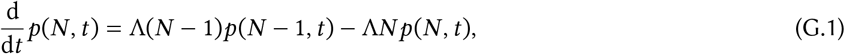

which yields a shifted negative binomial distribution of the form

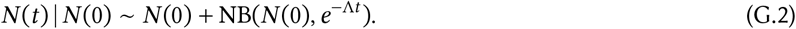

Starting from a single individual, one can take the limit *t* → ∞ to show that *Z* ~ Exp(1). As a result, E[log *Z*] = −γ ≈ −0.577, where γ is the Euler-Mascheroni constant, so

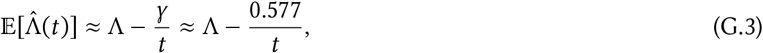

and the direct growth rate estimator has bias of order *O*(1/*t*). For the arithmetic mean (pooled) estimator in App. B, expanding *N* (*t*)/*N* (0) around its expected value for large *M* = *N* (0) shows that

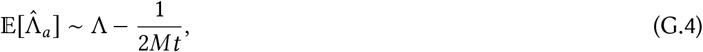

which remains biased. However, the fact that E[*Z*] = 1 in this example implies that the arithmetic mean estimator in App. B is asymptotically unbiased for large *M*, even for finite *t*. As discussed in App. A, this holds whenever the initial cell type is drawn from the steady-state distribution *π*_*i*_, which in this case is trivial.

Thinning this population with a constant rate *δ* turns this into a linear birth-death process described by the equation

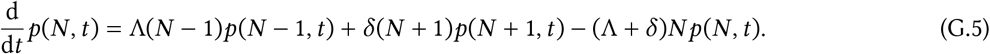

Now there is a nonzero chance of extinction, which at time *t* equals

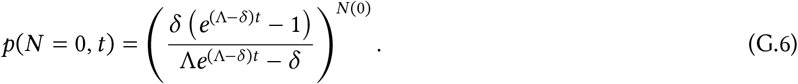

For *δ* ≥ Λ, this probability approaches 1 in the long run, so the population will almost surely become extinct. Furthermore, the probability of extinction decreases exponentially with the initial population size *N* (0): this is because each individual produces its own independent offspring, and the total population becomes extinct only when the populations founded by each of the initial individuals become extinct. For *δ* > Λ, the rate of extinction is *δ* − Λ < *δ*, so we can typically simulate a population for a few half-lives without risking extinction (depending on *N* (0)).

The strict chemostat is even easier to analyse than the full population, since all cells are statistically indistinguishable. Since all cells divide with rate Λ, birth events follow a Poisson process with rate *L*Λ. In particular, *B*(*t*) ~ Poi(Λ*Lt*), and using the probability generating function of the Poisson distribution we can directly verify that the population size estimator Eq. (16) is unbiased:

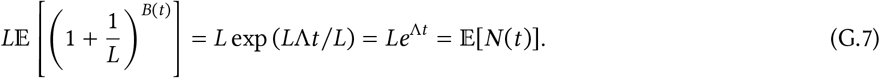

In contrast, the estimated growth rate would be

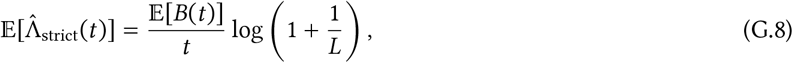

which for large *L* is approximately

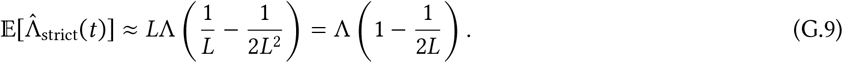

We therefore see that the finite-size bias is of order 1/*L*. In the single-lineage case,

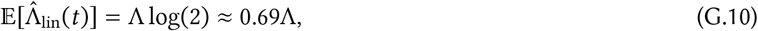

which has (relative) error 1 − log(2) ≈ 31%. The variance of the growth rate estimate is

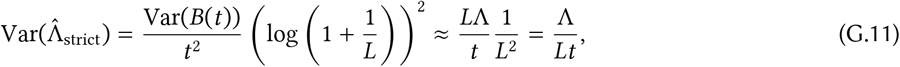

which is of order *O*(1/*Lt*), as expected.

We remark that the finite-time bias vanishes for the strict chemostat, as it does for the arithmetic mean estimator in the large-*M* limit. This is because the population is always at steady state, since the process is entirely memoryless. This does not hold for general branching processes.

### H Numerical details

We implemented the numerical models in this paper in Julia using <monospace>DifferentialEquations.j</monospace>l [96], <monospace>ModelingToolkit.jl</monospace> [97] and Catalyst.jl [98]; code can be found at https://github.com/kaandocal/chemostat_models. The algorithms discussed in this paper are available as the Julia package Chemostats.jl at https://github.com/kaandocal/Chemostats.jl.

The results in Fig. 1**d** were obtained by running 10 simulations using the strict chemostat for each data point. The linear birth model is that described in App. G, where each cell divides after an exponentially distributed amount of time with rate 1. The adder model was taken from [99]: each cell has volume *V* growing exponentially with rate 1, and a cell divides when its volume reaches its birth volume *V*_*b*_ plus a random division threshold Δ ~ Γ(5, 0.2), where 0.2 is the scale parameter. Upon division, the volume is partitioned at random between the daughter cells, with the fraction inherited by a daughter uniformly distributed on [0, 1]. Since the total volume of the population grows exponentially with rate 1, and since the average size per cell stays roughly constant, the number of cells must also grow with rate Λ = 1 [39].

### I Tetracycline model

We describe a cell using its volume *V* and intracellular tetracycline concentration [Tc], as well as the number R of TetR and A of TetA molecules. Cell volume grows as

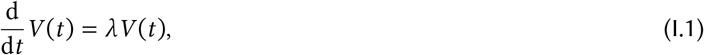

with instantaneous growth rate λ determined by Eq. (18). The intracellular tetracycline concentration follows Eq. (23), where [Tc]_ext_ is a fixed time-dependent input (described below). TetR and TetA are modelled as discrete variables, produced as inhomogeneous Poisson processes with rates Eq. (21) and Eq. (22), approximating O_1_ and O_2_ as described below. We do not model explicit degradation of TetR and TetA, only dilution due to cell growth and division.

Following [65], we derive the nutritional capacity κ_*n*_ from the maximal growth rate λ_max_ in the absence of tetracycline and TetA as

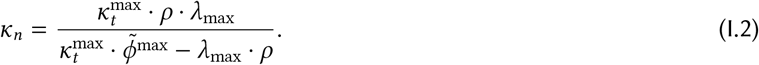

We assume that TetR molecules within a cell dimerise instantly, so that the number of dimers in a cell is given by

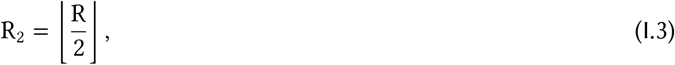

with 0 or 1 unpaired TetR molecules present at any given time. We model the number of TetR dimers bound to tetracycline via the reaction

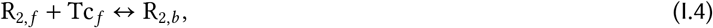

where R_2,*f*_ and R_2,*b*_ denote free and bound TetR dimers, respectively. Here Tc_*f*_ = Tc − R_2,*b*_ is the number of free tetracycline molecules, equal to *N*_*A*_ ⋅ *V* times the concentration, with *N*_*A*_ equal to Avogadro’s constant. Since tetracycline is much more abundant than TetR, we neglect tetracycline sequestration due to binding in Eqs. (20) and (23). As tetracycline binds and unbinds very quickly, we apply the quasiequilibrium approximation to Eq. (I.4) as in [65] to obtain

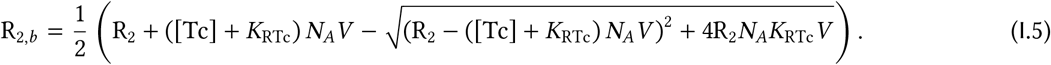

We assume that TetR dimers can bind independently to the operator sequences O_1_ and O_2_ with dissociation constants *K*_O1_ and *K*_O2_. For a fixed number of dimers, the joint probabilities that these two sites are bound or unbound become

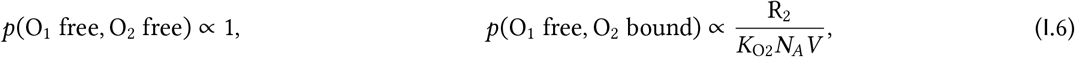

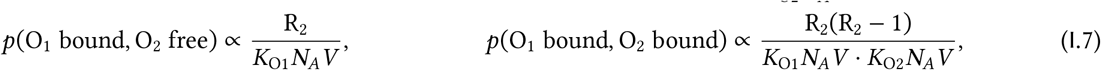

up to normalisation. We assume that TetR dimers bound to tetracycline do not inhibit transcription. Since TetR binding/unbinding events happen much faster than transcription, we again use the quasiequilibrium approximation to estimate the probability that the TetR transcription start site *P*_*R*2_ (Fig. 3) is active as

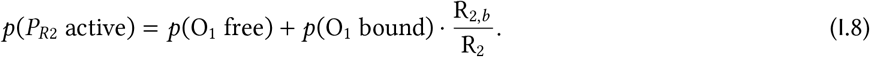

Similarly, we compute the probability that the transcription start sites *P*_*R*1_ and *P*_*A*_ are active as

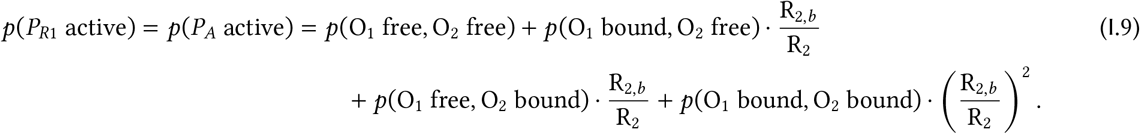

**Table S1.** Parameter values used for the tetracycline resistance model.

| Parameter | Value | Notes |
|---|---|---|
| *Cell growth* |  |  |
| *λ*<sub>max</sub> | log(2)/(45 min) | Adapted from [65]. |
| *V*<sub>*d*</sub> | 2 fL | Average bacterial cell volume is ≈ 1.4 fL [100]. |
| *ρ* | 0.76 | [66] |
| *φ̃*<sup>max</sup> | 0.48 | [66] |
| *κ*<sub>*t*</sub><sup>max</sup> | 4.5 h<sup>−1</sup> | [66] |
| *X*<sub>Tc</sub> | 1 µM | Adapted from [65]. |
| *X*<sub>A</sub> | 15 fL<sup>−1</sup> | See text. |
| *Tetracycline transport* |  |  |
| *D*<sub>Tc</sub> | 0.015 min<sup>−1</sup> | [65] |
| *α* | 2 min<sup>−1</sup> | See text. |
| *K*<sub>A</sub> | 10 µM | [65] |
| *Gene expression* |  |  |
| *k*<sub>R</sub> | 0.2 min<sup>−1</sup> fL<sup>−1</sup> | Rounded from [65]. |
| *k*<sub>A</sub> | 0.2 min<sup>−1</sup> fL<sup>−1</sup> | See text. |
| *K*<sub>RTc</sub> | 1 nM | [65] |
| *K*<sub>O1</sub> | 1 pM | Rounded from [62], depends on Na concentration [101]. |
| *K*<sub>O2</sub> | 0.5 pM | O<sub>2</sub> binds TetR with twice the affinity of O<sub>1</sub> [101]. |
| *Tetracycline treatment* |  |  |
| *C* | 100 µM | — |
| *τ*<sub>Tc</sub> | 100 min | — |
| *f*<sub>Tc</sub> | 20% | — |

This yields the effective production rates

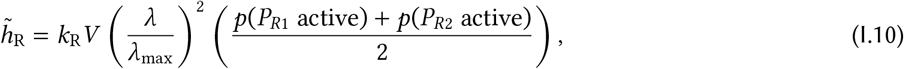

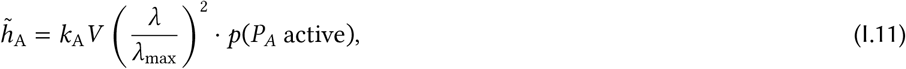

Cells divide once they reach a fixed division volume *V*_*d*_, and produce two daughter cells with exactly half their parent’s volume. TetA molecules are independently assigned at random to the daughters (symmetric binomial partitioning). We assume that each daughter inherits its parent’s tetracycline concentration. For TetR, we found that random partitioning of dimers resulted in daughter cells frequently inheriting no TetR due to low molecule numbers, which led to spurious production of TetA immediately following division. When there are at least two dimers in the mother cell, therefore, we assume that each daughter inherits one dimer, with the rest partitioned binomially. The single unpaired TetR molecule, when present, is inherited by a random daughter.

Since we were not able to find typical abundances of TetA in *E. coli*, we chose the transcription rate to be equal to that of TetR. For the export efficiency *α*, we chose a value that resulted in low intracellular tetracycline levels at 10 TetA molecules. For the inhibition constant *X*_A_, we chose a value corresponding to 15–30 molecules per cell, resulting in moderate growth inhibition at the concentrations we observed in our simulations (cf. Fig. 3**d**). We note that the system size parameter Ω rescales precisely these three parameters. Our simulations for Ω = 1 yield trajectories similar to those of [65].

We modelled extracellular tetracycline as a step function, with periodic tetracycline pulses at intervals of length *τ*_Tc_. Every pulse consists of a concentration *C* of tetracycline maintained for *f*_Tc_ ⋅ *τ*_Tc_ minutes, so that cells are exposed to tetracycline a fixed fraction *f*_Tc_ of the time (maintaining the same total dose for different intervals). For Fig. 3**c**, strictly periodic treatment intervals led to resonance with the (mostly deterministic) cell cycle. To avoid the resulting artefacts, we modelled the duration between pulses as a Gamma-distributed variable with mean *τ*_Tc_ and a coefficient of variation of 10%.

We simulated individual bacteria in Julia, solving the ordinary differential equations using the Bogacki-Shampine 3(2) method implemented in <monospace>DifferentialEquations.jl</monospace>. For the lax chemostat, we used an adaptation time of *τ*_*a*_ = 50 min with smoothing parameter *β* = 0.5. Backward lineages in Fig. 3**d** were obtained by simulating the lax chemostat with *L* = 100, and tracing the ancestry of a randomly sampled cell in the final population. Data for Fig. 3**e** and **f** was obtained using the lax chemostat with *L* = 100. All data points in Fig. 3 were averaged over 3 independent runs, simulated until convergence as determined by visual inspection. Where not otherwise noted, model parameters for all simulations are given in Table S1.

### J Chemotaxis model

Each cell is described by its volume *V*, its position (*x* and *y* coordinates), its angular orientation *θ*, and its methylation level *m*. The methylation level *m* evolves according to Eq. (26). By suitably shifting and rescaling *m*, we can assume that *ε*_0_ = 0 and *ε*_1_ = −1 in Eq. (28). The nutrient concentration ν(*t*) is measured at the current position of the cell. Cell volume grows as

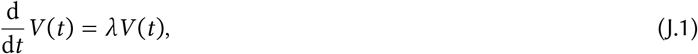

where the instantaneous growth rate λ is given in Eq. (30). Cells divide when they reach a fixed division volume *V*_*d*_. Each daughter cell inherits exactly half its parent’s volume, together with its parent’s position (*x, y*), methylation level *m* and, for simplicity, orientation *θ* (the orientation will be randomised by the next tumble).

During a run, the cell moves in the direction of *θ* with constant speed *v*_max_,

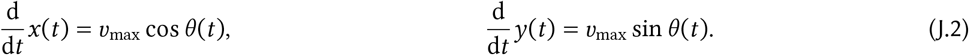

We model rotational diffusion using the stochastic differential equation

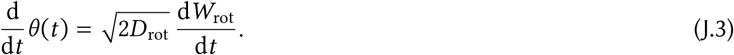

Here *D*_rot_ was chosen to match the mean square change in angle during runs in [69]. The cell switches from a run to a tumble stochastically with time-dependent rate Eq. (25). When a tumble occurs, the cell stops for a time *τ*_tum_ and changes its orientation by a random amount. In three dimensions, [69] reports a mean change of 69^°^, but since three-dimensional rotations cannot directly be projected to two dimensions, we sample a random increment 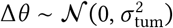.

We simulate cells in a two-dimensional environment of width *L*_*x*_ and height *L*_*y*_ with periodic boundary conditions (mathematically, a torus). Nutrients are added in patches following a homogeneous Poisson point process with intensity *I*_ν_, with each patch containing a fixed total amount ν_*p*_. Nutrients diffuse with diffusivity *D*_ν_, and are degraded exponentially with time constant *τ*_ν_. The nutrient concentration from a patch occurring at position (*x*_0_, *y*_0_) at time *t*_0_ is described by the initial value problem

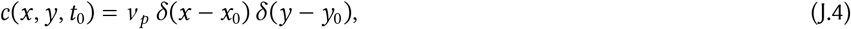

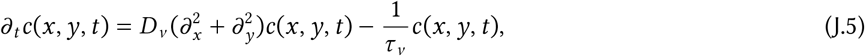

for *t* > *t*_0_, with periodic boundary conditions

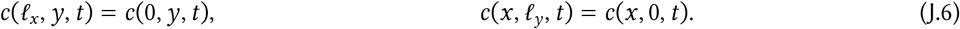

We can solve the diffusion equation analytically as

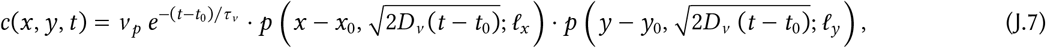

where

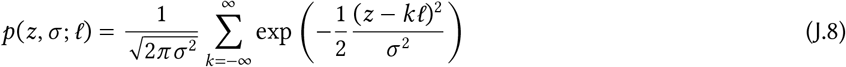

is the probability density of the wrapped normal distribution with standard deviation *σ* and periodicity *L*. The total nutrient concentration ν is obtained by summing over all patches.

**Table S2.**
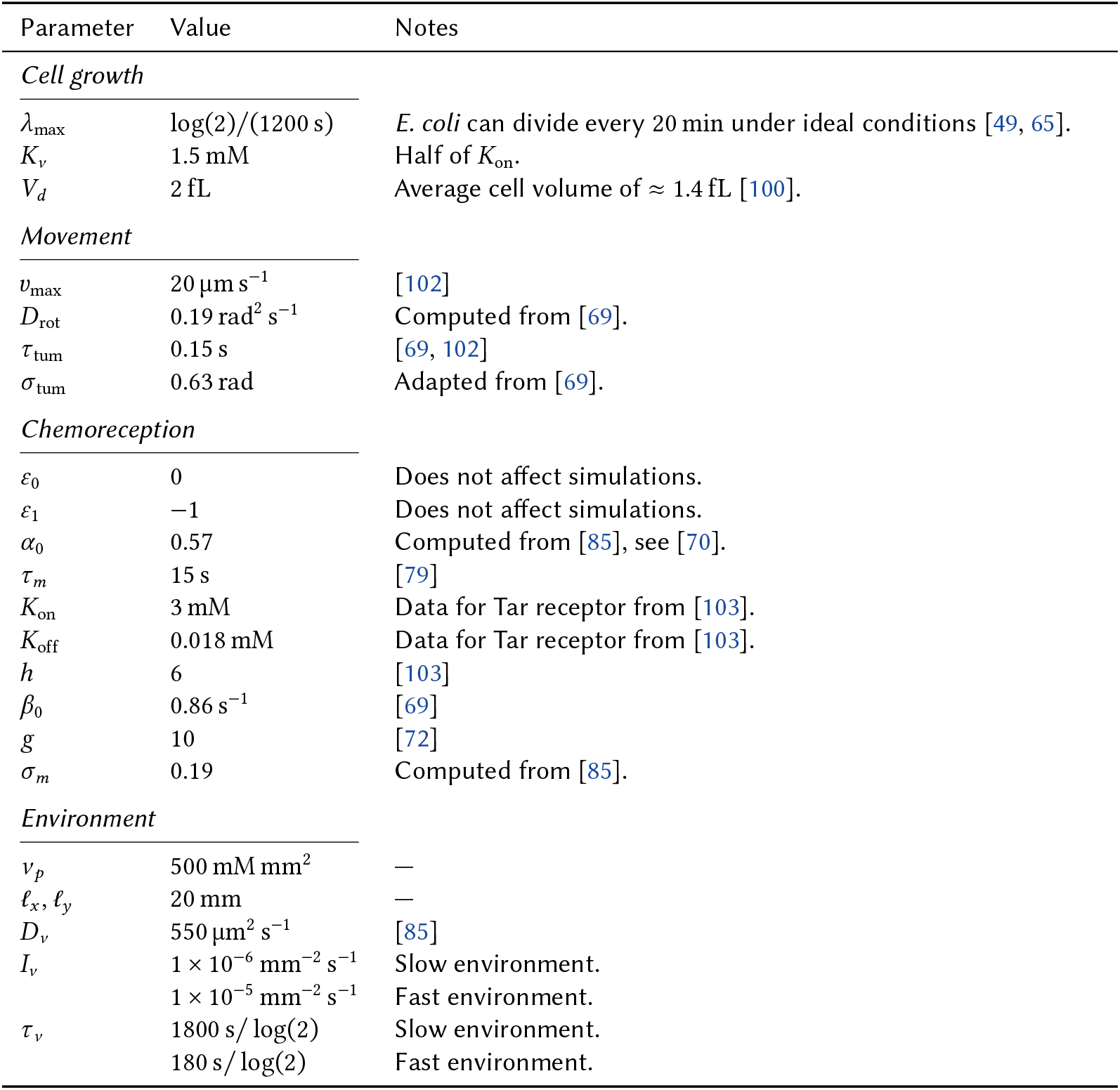
Parameter values used for the chemotaxis model.

| Parameter | Value | Notes |
|---|---|---|
| *Cell growth* |  |  |
| *λ*<sub>max</sub> | log(2)/(1200 s) | *E. coli* can divide every 20 min under ideal conditions [49, 65]. |
| *K*<sub>*ν*</sub> | 1.5 mM | Half of *K*<sub>on</sub>. |
| *V*<sub>*d*</sub> | 2 fL | Average cell volume of ≈ 1.4 fL [100]. |
| *Movement* |  |  |
| *v*<sub>max</sub> | 20 µm s<sup>−1</sup> | [102] |
| *D*<sub>rot</sub> | 0.19 rad<sup>2</sup> s<sup>−1</sup> | Computed from [69]. |
| *τ*<sub>tum</sub> | 0.15 s | [69, 102] |
| *σ*<sub>tum</sub> | 0.63 rad | Adapted from [69]. |
| *Chemoreception* |  |  |
| *ε*<sub>0</sub> | 0 | Does not affect simulations. |
| *ε*<sub>1</sub> | −1 | Does not affect simulations. |
| *α*<sub>0</sub> | 0.57 | Computed from [85], see [70]. |
| *τ*<sub>*m*</sub> | 15 s | [79] |
| *K*<sub>on</sub> | 3 mM | Data for Tar receptor from [103]. |
| *K*<sub>off</sub> | 0.018 mM | Data for Tar receptor from [103]. |
| *h* | 6 | [103] |
| *β*<sub>0</sub> | 0.86 s<sup>−1</sup> | [69] |
| *g* | 10 | [72] |
| *σ*<sub>*m*</sub> | 0.19 | Computed from [85]. |
| *Environment* |  |  |
| *ν*<sub>*p*</sub> | 500 mM mm<sup>2</sup> | — |
| *ℓ*<sub>*x*</sub>, *ℓ*<sub>*y*</sub> | 20 mm | — |
| *D*<sub>*ν*</sub> | 550 µm<sup>2</sup> s<sup>−1</sup> | [85] |
| *I*<sub>*ν*</sub> | 1 × 10<sup>−6</sup> mm<sup>−2</sup> s<sup>−1</sup> | Slow environment. |
|  | 1 × 10<sup>−5</sup> mm<sup>−2</sup> s<sup>−1</sup> | Fast environment. |
| *τ*<sub>*ν*</sub> | 1800 s/ log(2) | Slow environment. |
|  | 180 s/ log(2) | Fast environment. |

We simulated individual bacteria in Julia, solving the differential equations using the Bogacki-Shampine 3(2) method implemented in `DifferentialEquations.jl`. We implemented the rotation and signalling noise in Eqs. (26) and (J.3) by adding Gaussian noise at intervals of length 0.1 s. For the lax chemostat, we used an adaptation time of *τ*_*a*_ = 1000 s with smoothing parameter *β* = 0.5, except for *L* = 50, where we used an adaptation time of *τ*_*a*_ = 500 s. All data points in Fig. 4 were averaged over 3 independent runs, simulated until convergence as determined by visual inspection. Where not otherwise noted, model parameters for all simulations are given in Table S2.

